# Mice alternate between discrete strategies during perceptual decision-making

**DOI:** 10.1101/2020.10.19.346353

**Authors:** Zoe C. Ashwood, Nicholas A. Roy, Iris R. Stone, The International Brain Laboratory, Anne E. Urai, Anne K. Churchland, Alexandre Pouget, Jonathan W. Pillow

## Abstract

Classical models of perceptual decision-making assume that subjects use a single, consistent strategy to form decisions, or that strategies evolve slowly over time. Here we present new analyses suggesting that this common view is incorrect. We analyzed data from mouse and human decision-making experiments and found that choice behavior relies on an interplay between multiple interleaved strategies. These strategies, characterized by states in a hidden Markov model, persist for tens to hundreds of trials before switching, and may alternate multiple times within a session. The identified mouse decision-making strategies were highly consistent across individuals and comprised a single “engaged” state, in which decisions relied heavily on the sensory stimulus, and several biased states in which errors frequently occurred. These results provide a powerful alternate explanation for “lapses” often observed in rodent psychophysical experiments, and suggest that standard measures of performance mask the presence of dramatic changes in strategy across trials.

## 1 Introduction

Understanding the computations performed in the brain will require a comprehensive characterization of behavior [1–3]. This realization has fueled a recent surge in methods devoted to the measurement, quantification, and modeling of natural behaviors [4–8]. Historically, studies of perceptual decision-making behavior have tended to rely on models derived from signal detection theory (SDT) [9, 10] or evidence accumulation [11–13]. More recently, approaches based on reinforcement learning have also been used to model the effects of context, reward, and trial-history on perceptual decision-making behavior [14–18]. In all cases, however, these approaches describe decision-making in terms of a single strategy that does not change abruptly across trials or sessions.

One puzzling aspect of sensory decision-making behavior is the presence of so-called “lapses”, in which an observer makes an error despite the availability of strong sensory evidence. The term itself suggests an error that arises from a momentary lapse in attention or memory, as opposed to an inability to perceive the sensory stimulus. Lapses arise in all species, but are surprisingly frequent in rodent experiments, where lapses can comprise up to 10-20% of all trials [19–21].

The standard approach for modeling lapses involves augmenting the classic psychometric curve with a “lapse parameter”, which characterizes the probability that the observer simply ignores the stimulus on any given trial [22–24]. This model can be conceived as a mixture model [25] in which, on every trial, the animal flips a biased coin to determine whether or not to pay attention to the stimulus when making its choice. Previous literature has offered a variety of explanations for lapses, including inattention, motor error, and incomplete knowledge of the task [22, 24, 26], and recent work has argued that they reflect an active process of uncertainty-guided exploration [17]. However, a common thread to these explanations is that lapses arise independently across trials, in a way that does not depend on the time course of other lapses.

Here we show that lapses do not arise independently, but depend heavily on latent states that underlie decision-making behavior. We use a modeling framework based on hidden Markov models (HMMs) to show that mice rely on discrete decision-making strategies that persist for tens to hundreds of trials. While the classic lapse model corresponds to a special case in our framework, the model that best described the empirical choice data of real mice had one state corresponding to an “engaged” strategy, in which the animal’s choices were strongly influenced by the sensory stimulus, and other states that corresponded to biased or weakly stimulus-dependent strategies. Our analyses show that lapses arise primarily during long sequences of trials when the animal is in a biased or disengaged state. Conversely, we found that animals with high apparent lapse rates may nevertheless be capable of high-accuracy performance for extended blocks of trials. We applied our modeling framework to datasets from two different mouse decision-making experiments [19, 20], and to one dataset from a decision-making experiment in humans [27]. Taken together, these results shed substantial new light on the factors governing sensory decision-making, and provide a powerful set of tools for identifying hidden states in behavioral data.

## 2 Results

### 2.1 The classic lapse model for sensory decision-making

A common approach for analyzing data from two-choice perceptual decision-making experiments involves the psychometric curve, which describes the probability that the animal chooses one option (e.g., “rightward”) as a function of the stimulus value [22–24]. The psychometric curve is commonly parameterized as a sigmoidal function that depends on a linear function of the stimulus plus an offset or bias. This sigmoid rises from a minimum value of *γ*_*r*_ to a maximal value of 1 − *γ*_*l*_, where *γ*_*r*_ and *γ*_*l*_ denote “lapse” parameters, which describe the probability of of making a rightward or leftward choice independent of the stimulus value. Thus, the probability of a rightward choice is always at *least γ*_*r*_, and it cannot be greater than 1 − *γ*_*l*_. In what follows, we will refer to this as the “classic lapse model of choice behavior”, which can be written:

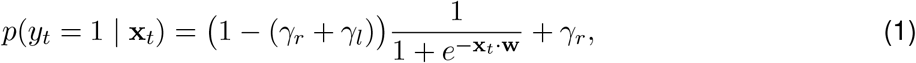

where *y*_*t*_ ∈ {0, 1} represents the choice (left or right) that an animal makes at trial *t*, **x**_*t*_ ∈ ℝ^*M*^ is a vector of covariates, and **w** ∈ ℝ^*M*^ is a vector of weights that describes how much each covariate influences the animal’s choice. Note that **x**_*t*_ includes both the stimulus and a constant ‘1’ element to capture the bias or offset, but it may also include other covariates that empirically influence choice, such as previous choices, stimuli, and rewards [28–30].

Although the classic lapse model can be viewed as defining a particular sigmoid-shaped curve relating the stimulus strength to behavior (Fig. 1c), it can equally be viewed as a mixture model [25]. In this interpretation, we regard the animal as having an internal state *z*_*t*_ that takes on one of two different values on each trial, namely “engaged” or “lapse”. If the animal is engaged, it makes its choice according to the classic sigmoid curve (which saturates at 0 and 1). If lapsing, it ignores the stimulus and makes its choice based only on the relative probabilities of a left and right lapse. Mathematically, this can be written:

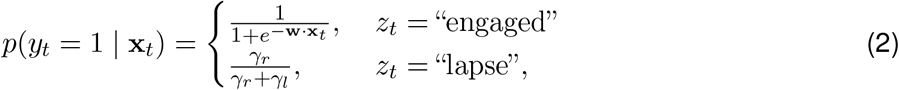

where *p*(*z*_*t*_ = “lapse”) = (*γ*_*r*_ + *γ*_*l*_) and *p*(*z*_*t*_ = “engaged”) = 1 − (*γ*_*r*_ + *γ*_*l*_). In this interpretation, the animal flips a biased coin, with fixed probability (*γ*_*r*_ + *γ*_*l*_), on each trial and then adopts one of two strategies based on the outcome—a strategy that depends on the stimulus vs. one that ignores it. Note that the animal can make a correct choice in the lapse state through pure luck; we use “lapse” here simply to indicate that the animal is not relying on the stimulus when making its decision.

**Figure 1:**
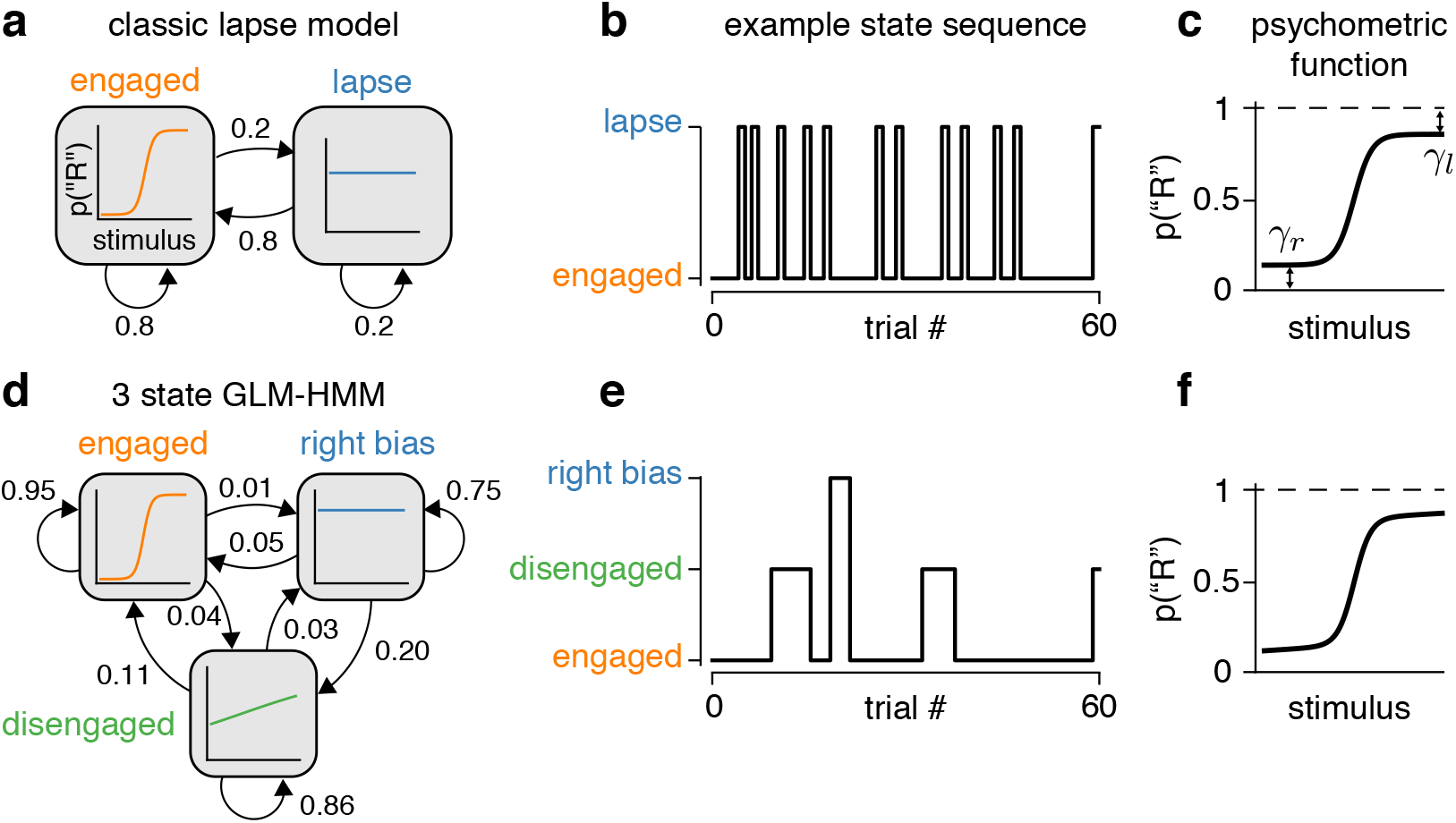
The GLM-HMM generalizes the classic lapse model. **(a)** The classic lapse model formulated as a 2-state GLM-HMM. Each box represents a generalized linear model (GLM) describing how the probability of a binary choice depends on the stimulus in the corresponding state: “engaged” (left) or “lapse” (right). Arrows between boxes indicate the transition probabilities between states. Note that the probability of switching to the engaged state at the next trial is always 0.8 (it is independent of state at the current trial) and similarly there is always a 0.2 probability of entering the lapse state on each trial.**(b)** An example (simulated) sequence for the animal’s internal state when the transitions between states are governed by the probabilities in (a). Notice that the lapse state tends to last for only a single trial at a time. **(c)** Psychometric function arising from the model shown in (a), depicting the probability of a rightward choice as a function of the stimulus. The parameters *γ*_*r*_ and *γ*_*l*_ denote the probability of a rightward and leftward lapse, respectively. As specified by the transition probabilities in (a), the total lapse probability for this model is *γ*_*r*_ + *γ*_*l*_ = 0.2. **(d**) Example 3-state GLM-HMM, with three different GLMs corresponding to different decision-making strategies (labeled “engaged”, “disengaged” and “right biased”). Note: these are just *example* states for the 3-state GLM-HMM. In reality, we will *learn* the states that best describe each animal’s choice data using the process described in Section 4.1. The high self-transition probabilities of 0.95, 0.86 and 0.75 ensure that these states typically persist for many trials in a row. **(e)** An example sequence for the animal’s internal state sampled from the GLM-HMM shown in (d). **(f)** The psychometric curve arising from the model shown in (d), which corresponds to a weighted average of the psychometric functions associated with each state. Note that although the decision-making models shown in (a) and (d) are vastly different, the resulting psychometric curves are nearly identical, meaning that the psychometric curve alone cannot be used to distinguish them.

Viewing the classic lapse model as a mixture model highlights some of its limitations. First, it assumes that animals switch between only two decision-making strategies. Second, it assumes that lapses occur independently in time, according to an independent Bernoulli random variable on each trial. Finally, the model assumes that choices in the “lapse” state are fully independent of the stimulus, neglecting the possibility that they are still weakly stimulus dependent [24], or are influenced by other covariates such as reward or choice history [28, 29]. These limitations motivate us to consider a more general family of models, which includes the classic lapse model as a special case.

### 2.2 A model for decision-making with multiple strategies

Recognizing the limitations of the classic lapse model, we propose to analyze perceptual decision-making behavior using a framework based on Hidden Markov models (HMMs) with Bernoulli generalized linear model (GLM) observations [6, 31]. The resulting “GLM-HMM” framework, also known as an input-output HMM [32], allows for an arbitrary number of states, which can persist for an extended number of trials and exhibit different dependencies on the stimulus and other covariates.

A GLM-HMM has two basic pieces: an HMM governing the distribution over latent states, and a set of state-specific GLMs, specifying the decision-making strategy employed in each state (see Fig. 1). For a GLM-HMM with *K* latent states, the HMM has a *K* × *K* transition matrix *A* specifying the probability of transitioning from any state to any other,

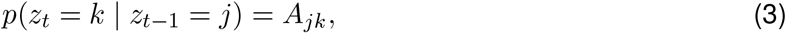

where *z*_*t*−1_ and *z*_*t*_ indicate the latent state at trials *t* − 1 and *t*, respectively. The “Markov” property of the HMM is that the state on any trial depends only on the state from the previous trial, and the “hidden” property refers to the fact that states are latent or hidden from external observers. For completeness, the HMM also has a distribution over initial states, given by a *K*-element vector ***π*** whose elements sum to 1, giving *p*(*z*_1_ = *k*) = ***π***_*k*_.

To describe the state-dependent mapping from inputs to decisions, the GLM-HMM contains *K* independent Bernoulli GLMs, each defined by a weight vector specifying how inputs are integrated in that particular state. The probability of a rightward choice (*y*_*t*_ = 1) given the input vector **x**_*t*_ and the latent state *z*_*t*_ is given by

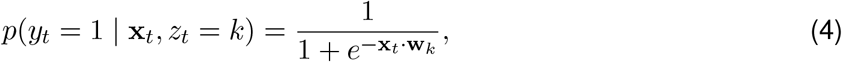

where **w**_*k*_ ∈ ℝ^*M*^ denotes the GLM weights for latent state *k* ∈ {1, …, *K*}. The full set of parameters for a GLM-HMM, 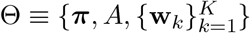, are learned directly from each animal’s choice data via the EM algorithm described in Section 4.1.

It is worth noting that the classic lapse model described in Eq. 1 and Eq. 2 corresponds to a restricted 2-state GLM-HMM. If we consider state 1 to be “engaged” and state 2 to be the “lapse” state, then the state-1 GLM has weights **w**_1_ = **w**, and the state-2 GLM has all weights set to 0 except the bias weight, which is equal to − log(*γ*_*l*_*/γ*_*r*_). The transition matrix has identical rows, with probability 1 − (*γ*_*r*_ + *γ*_*l*_) of going into state 1 and probability (*γ*_*r*_ + *γ*_*l*_) of going into state 2 at the next trial, regardless of the current state. This ensures that the probability of a lapse on any given trial is stimulus-independent and does not depend on the previous trial’s state. Fig. 1a-c shows an illustration of the classic lapse model formulated as a 2-state GLM-HMM.

However, there is no general reason to limit our analyses to this restricted form of the GLM-HMM. By allowing the model to have more than two states, multiple states with non-zero stimulus weights, and transition probabilities that depend on the current state, we obtain a model family with a far richer set of dynamic decision-making behaviors. Fig. 1d shows an example GLM-HMM with three latent states, all of which have high probability of persisting for multiple trials. Intriguingly, the psychometric curve arising from this model (Fig. 1f) is indistinguishable from that of the classic lapse model. Thus, multiple generative processes can result in identical psychometric curves, and we must look beyond the psychometric curve if we want to gain insight into the dynamics of decision-making across trials.

### 2.3 Mice switch between multiple strategies

To examine whether animals employ multiple strategies during decision-making, we fit the GLM-HMM to behavioral data from two binary perceptual decision-making tasks (see Methods 4.1). First, we fit the GLM-HMM to choice data from 37 mice performing a visual detection decision-making task developed in [33] and adopted by the International Brain Laboratory (IBL) [19]. During the task, a sinusoidal grating with contrast between 0 and 100% appeared either on the left or right side of the screen (Fig. 2a). The mouse had to indicate this side by turning a wheel. If the mouse turned the wheel in the correct direction, it received a water reward; if incorrect, it received a noise burst and an additional 1-second timeout. During the first 90 trials of each session, the stimulus appeared randomly on the left or right side of the screen with probability 0.5. Subsequent trials were generated in blocks in which the stimulus appeared on one side with probability 0.8, alternating randomly every 20-100 trials. We analyzed data from animals with at least 3000 trials of data (across multiple sessions) after they had successfully learned the task (see Fig. S1, Fig. S2). For each animal, we considered only the data from the first 90 trials of each session, when the stimulus was equally likely to appear on the left or right of the screen.

**Figure 2:**
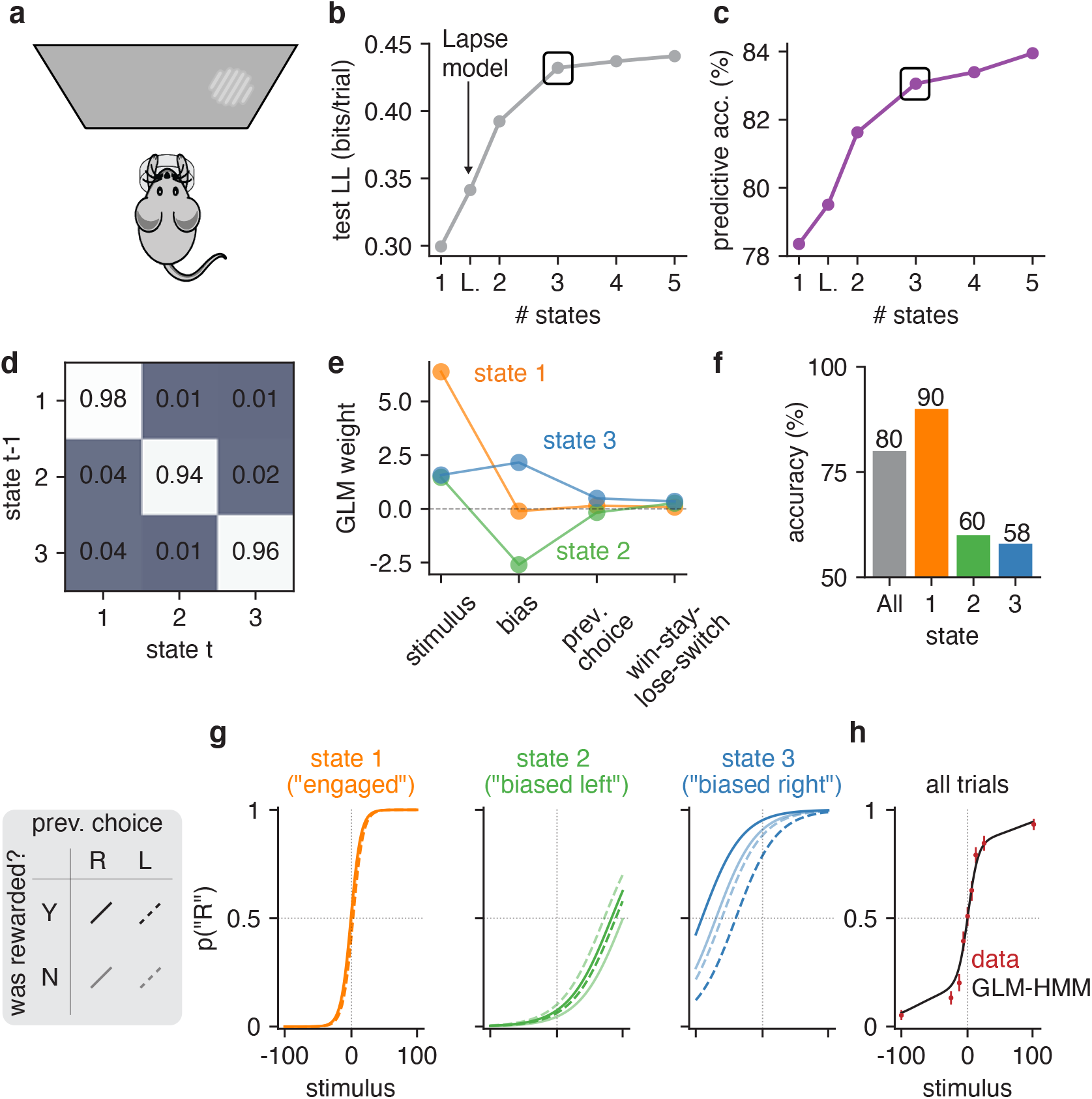
GLM-HMM analysis of choice behavior of an example IBL mouse. **(a)** Schematic for visual decision-making task, which involved turning a wheel to indicate whether a Gabor patch stimulus appeared on the left or right side of the screen [19, 33]. **(b)** Model comparison between GLM-HMMs with different numbers of latent states, as well as classic lapse model (labeled ‘L’) using 5-fold cross-validation. Test set log-likelihood is reported in units of bits/trial and was computed relative to a ‘null’ Bernoulli coin flip model (see Section 4.2.2). A black rectangle highlights the log-likelihood for the 3-state model, which we used for all subsequent analyses. **(c)** Test set predictive accuracy for each model, indicating the percent of held-out trials where the model successfully predicted the mouse’s true choice. **(d)** Inferred transition matrix for best-fitting 3 state GLM-HMM for this mouse. Large entries along the diagonal indicate a high probability of remaining in the same state. **(e)** Inferred GLM weights for the 3-state model. State 1 weights have a large weight on the stimulus, indicating an “engaged” or high-accuracy state. In states 2 and 3, the stimulus weight is small, and the bias weights give rise to large leftward (state 2) and rightward (state 3) biases. **(f)** Overall accuracy of this mouse (grey), and accuracy for each of the three states. **(g)** Psychometric curve for each state, conditioned on previous reward and previous choice. **(h)** The standard psychometric curve for all choice data from this mouse, which can be seen to arise from the mixture of the three per-state curves shown in panel g. Using the fit GLM-HMM parameters for this animal and the true sequence of stimuli presented to the mouse, we generated a time series with the same number of trials as those that the example mouse had in its dataset. At each trial, regardless of the true stimulus presented, we calculated *p*_*t*_(“*R*”) for each of the 9 possible stimuli by averaging the per-state psychometric curves of panel g and weighting by the appropriate row in the transition matrix (depending on the sampled latent state at the previous trial). Finally, we averaged the per-trial psychometric curves across all trials to obtain the curve that is shown in black, while the empirical choice data of the mouse are shown in red, as are 95% confidence intervals (n between 530 and 601, depending on stimulus value).

We modeled the animals’ decision-making strategies using a GLM-HMM with four inputs: (1) the (signed) stimulus contrast, where positive values indicate a right-side grating and negative values indicate a left-side grating; (2) a constant offset or bias; (3) the animal’s choice on the previous trial; and (4) the stimulus side on the previous trial. A large weight on the animal’s previous choice gives rise to a strategy known as “perserveration” in which the animal makes the same choice many times in a row, regardless of whether it receives a reward. A large weight on the previous stimulus side, which we refer to as the “win-stay, lose-switch” regressor, gives rise to the well-known strategy in which the animal repeats a choice if it was rewarded, and switches choices if it was not. Note that for the IBL task in question, bias and trial-history dependencies were sub-optimal, meaning that the maximal-reward strategy was to have a large weight on the stimulus and zero weights on the other three inputs.

To determine the number of different strategies underlying decision-making behavior, we fit GLM-HMMs with varying numbers of latent states. Note that the 1-state model is simply a standard Bernoulli GLM, while the classic lapse model (Eq. 1, Eq. 2) is the constrained 2-state model mentioned previously. We found that a 3-state GLM-HMM substantially outperformed models with fewer states, including the classic lapse model. The states of the fitted model were readily interpretable, and tended to persist for many trials in a row.

Figures 2 and 3 show results for an example mouse. For this animal, the multi-state GLM-HMM out-performed both the standard (1-state) GLM and the classic lapse model, both in test log-likelihood and percent correct, with the improvement approximately leveling off at 3 latent states (Fig. 2b-c). Because the test set for this mouse contained 900 trials, test set log-likelihood increases of 0.13 bits/trial and 0.09 bits/trial for the 3-state model over the 1-state model and classic lapse models, respectively, meant that the data were (2^0.13^)^900^ ≈ 1.7 × 10^35^ and (2^0.09^)^900^ ≈ 2.4 × 10^24^ times more likely under the 3-state model. Note that while the number of model parameters increased with the number of states in the GLM-HMM (and a two state GLM-HMM has more parameters than both the classic lapse model or single state GLM), the log-likelihood of the test dataset was not guaranteed to increase with more states [34]. Indeed, as the number of states increased, the GLM-HMM could have started to overfit the training dataset causing it to fit the test dataset poorly (see Fig. ED1).

**Figure 3:**
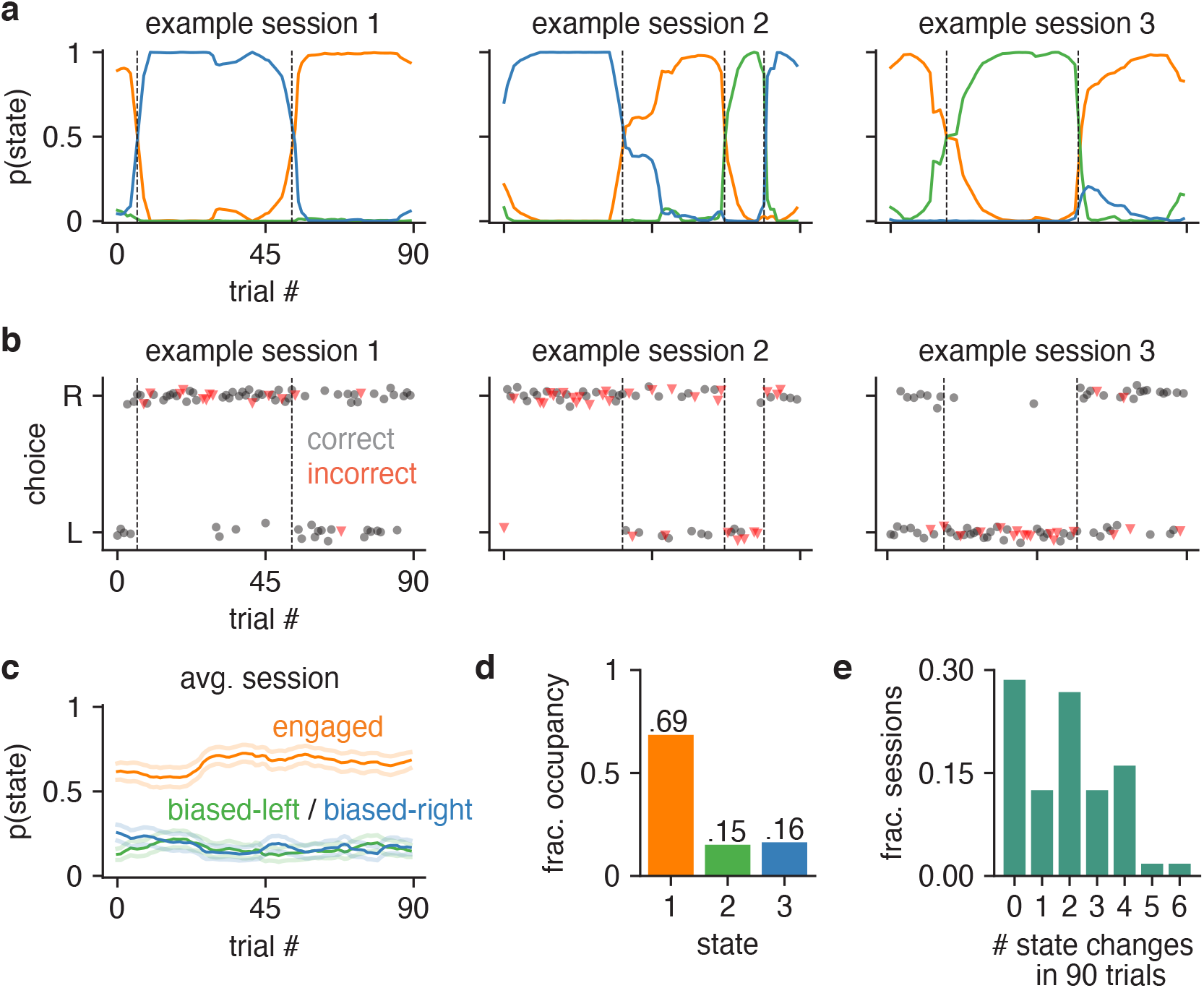
Inferred state dynamics for example IBL mouse. **(a)** Posterior state probabilities for three example sessions, revealing high levels of certainty about the mouse’s internal state, and showing that states typically persisted for many trials in a row. **(b)** Raw behavioral data corresponding to these three sessions. Each dot corresponds to a single trial; the x-position indicates the trial within the session, while the y-position indicates if the mouse went rightward or leftward on the trial (y-positions are slightly jittered for visibility). The dot’s color and shape indicates if the trial was an error or correct trial (red triangles correspond to errors, while black circles represents correct choices). All trials except 0% contrast trials (for which the correct answer is determined randomly) are shown here. The dashed vertical lines correspond to the location of state changes (obtained using the posterior probabilities shown in a). **(c)** Average trajectories of state probabilities within a session, computed over 56 sessions. Error bars indicate ±1 standard error of the mean. **(d)** Fractional occupancies for the three states across all trials. For this analysis, we assigned each trial to its most likely state and then counted the fraction of trials assigned to each state. Overall, the mouse spent 69% of all trials in the engaged state, and 31% of trials in one of the two biased states. **(e)** Histogram showing the number of inferred state changes per session, for all 56 sessions of data for this mouse. Only 29% of sessions involved the mouse persisting in a single state for the entire 90-trial session.

The transition matrix for the fitted 3-state model describes the transition probabilities between three different states, each of which corresponds to a different decision-making strategy (Fig. 2d). Large entries along the diagonal of this matrix, ranging between 0.94 and 0.98, indicate a high probability of remaining in the same state for multiple trials. The other set of inferred parameters were the GLM weights, which define how the animal makes decisions in each state (Fig. 2e). One of these GLMs (“state 1”) had a large weight on the stimulus and negligible weights on other inputs, giving rise to high-accuracy performance on the task (Fig. 2f). The other two GLMs (“state 2” and “state 3”), by comparison, had small smaller weights on the stimulus, and relatively large bias weights.

We can visualize the decision-making strategies associated with these states by plotting the corresponding psychometric curves (Fig. 2g), which show the probability of a rightward choice as a function of the stimulus, conditioned on both previous choice and reward. The steep curve observed in state 1, which corresponds to the mouse achieving near-perfect performance on high-contrast stimuli, led us to adopt the name ‘engaged’ to describe this state. By comparison, the psychometric curves for states 2 and 3 reflected large leftward and rightward biases, respectively. They also had relatively large dependence on previous choice and reward, as indicated by the gap between solid and dashed lines. While this mouse had an overall accuracy of 80%, it achieved 90% accuracy in the engaged state, compared to only 60% and 58% accuracy in the two biased states (Fig. 2f).

To gain insight into the temporal structure of decision-making behavior, we used the fitted 3-state model to compute the posterior probability over the mouse’s latent state across all trials (Fig. 3). The resulting state trajectories reflect our posterior beliefs about the animal’s internal state on every trial, given the entire sequence of observed inputs and choices during a session (See Methods, Sec. 4.1.3).

Contrary to the predictions of the classic lapse model, states tended to persist for many trials in a row. Remarkably, we found that the most probable state often had probability close to 1, indicating a high degree of confidence about the mouse’s internal state given the observed data.

To quantify state occupancies, we assigned each trial to its most probable state, and found that this example mouse spent approximately 69% of all trials in the engaged state (out of 5040 total trials over 56 sessions), compared to 15% and 16% of trials in the biased-left and biased-rightward states (Fig. 3c). Moreover, the mouse changed state at least once within a session in roughly 71% of all 90-trial sessions, and changed multiple times in 59% of sessions (Fig. 3d). This rules out the possibility that the states merely reflect the use of different strategies on different days. Rather, the mouse tended to remain in an engaged, high-performance state for tens of trials at a time, with lapses arising predominantly during interludes when it adopted a left-biased or right-biased strategy for multiple trials in a row. The multi-state GLM-HMM thus provides a very different portrait of mouse decision-making behavior than the basic GLM or lapse model.

### 2.4 State-based strategies are consistent across mice

To assess the generality of these findings, we fit the GLM-HMM to the choice data from 37 mice in the IBL dataset [19] (Fig. 4). We found that the results shown for the example animal considered above were broadly consistent across animals. Specifically, we found that the 3-state GLM-HMM strongly outperformed the basic GLM and classic lapse model in cross-validation for all 37 mice (Fig. 4a and Fig. S3). On average, it predicted mouse choices with 4.2% higher accuracy than the basic GLM (which had an average prediction accuracy of 78%), and 2.8% higher accuracy than the classic lapse model (Fig. 4b). Furthermore, for one animal the improvement in prediction accuracy for the 3 state GLM-HMM was as high as 12% relative to the basic GLM, and 7% relative to the classic lapse model.

**Figure 4:**
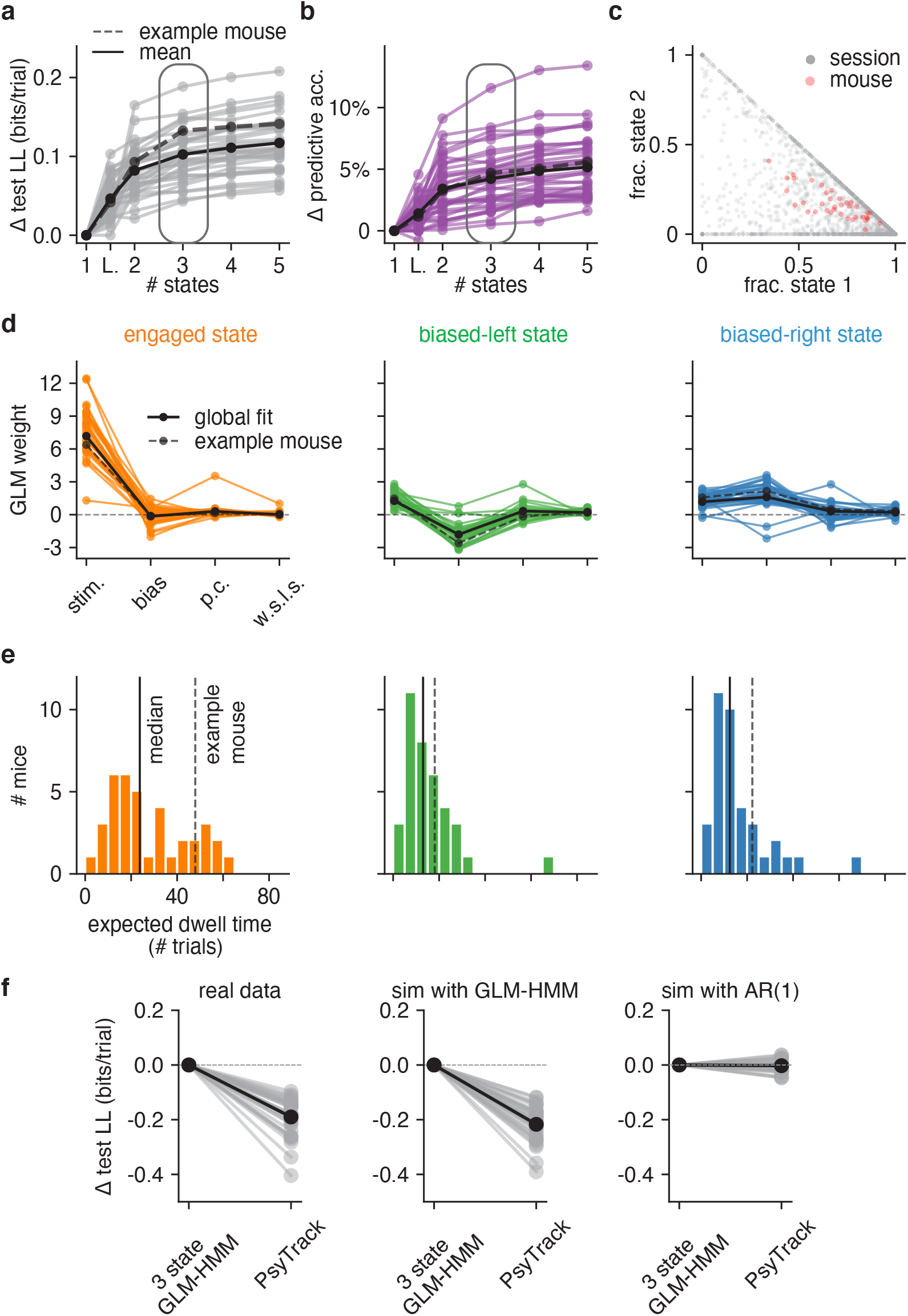
Model fits to full population of 37 IBL mice. **(a)** Change in test log-likelihood as a function of number of states relative to a (1-state) GLM, for each mouse in the population. The classic lapse model, a restricted form of the 2-state model, is labeled ‘L’. Each trace represents a single mouse. Solid black indicates the mean across animals, and the dashed line indicates the example mouse from Figs. 2 and 3. The rounded rectangle highlights performance of the 3-state model, which we selected for further analyses. **(b)** Change in predictive accuracy relative to a basic GLM for each mouse, indicating the percentage improvement in predicting choice. **(c)** Grey dots correspond to 2017 individual sessions across all mice, indicating the fraction of trials spent in states 1 (engaged) and 2 (biased left). Points at the vertices (1,0), (0,1), or (0,0) indicate sessions with no state changes, while points along the sides of the triangle indicate sessions that involved only 2 of the 3 states. Red dots correspond the same fractional occupancies for each of the 37 mice, revealing that the engaged state predominated, but that all mice spent time in all 3 states. **(d)** Inferred GLM weights for each mouse, for each of the three states in the 3-state model. The solid black curve represents a global fit using pooled data from all mice (see Algorithm 1); the dashed line is the example mouse from Fig. 2 and Fig. 3. **(e)** Histogram of expected dwell times across animals in each of the three states, calculated from the inferred transition matrix for each mouse. (**f**) Mice have discrete—not continuous—decision-making states. Left: Cross-validation performance of the 3 state GLM-HMM compared to PsyTrack [35, 36] for all 37 mice studied (each individual line is a separate mouse; black is the mean across animals). Middle: As a sanity check, we simulated datasets from a 3 state GLM-HMM with the parameters for each simulation chosen as the best fitting parameters for a single mouse. We then fit the simulated data both with PsyTrack and with the 3 state GLM-HMM in order to check that the 3 state GLM-HMM best described the data. Right: We did the opposite and fit PsyTrack to the animals’ data and then generated data according to an AR(1) model with parameters specified using the PsyTrack fits (see section 4.3 for full details). By performing cross-validation on the simulated data, we confirmed that we could use model comparison to distinguish between discrete and continuous decision-making behavior in choice data.

Although performance continued to improve slightly with four and even five latent states, we will focus our analyses on the 3-state model for reasons of simplicity and interpretability. Fig. S4 provides a full analysis of 4-state model, showing that it tended to divide the engaged state from the 3-state model into two “sub-states” that differed slightly in bias.

Fits of the 3-state GLM-HMM exhibited remarkable consistency across mice, with a majority exhibiting states that could be classified as “engaged”, “biased-left”, and “biased-right” (Fig. 4d). (See Methods section 4.1.5 for details about the alignment of states across mice). While we plot inferred transition matrices for all 37 mice in Fig. S5, here we used the diagonal elements of each matrix to compute an expected dwell time for each animal in each state (Fig. 4e). This revealed a median dwell time, across animals, of 24 trials for the engaged state, versus 13 and 12 trials for the biased-left and biased-right states, respectively. This description of behavior departs dramatically from the assumptions of the classic lapse model: for the classic lapse model and a lapse rate of 20%, the expected dwell time in the lapse state is just 1/(1-0.2) = 1.25 trials. We analyzed the distribution of state dwell times inferred from data and found they were well approximated by a geometric distribution, matching the theoretical distribution of data sampled from a Hidden Markov Model (Fig. ED2 and Fig. S6).

We also examined the fraction of trials per session that mice spent in each of the three states (Fig. 4c). To do so, we used the fitted model parameters to compute the posterior probabilities over state, and assigned each trial to its most likely state. The resulting “fractional occupancies” revealed that the median mouse spent 69% of its time in the engaged state, with the best mice exceeding 90% engagement. Moreover, the majority of sessions (83% of 2017 sessions across all animals) involved a switch between two or more states; in only 17% of sessions did a mouse remain in the same state for an entire 90-trial session.

To further quantify the model’s ability to capture temporal structure of choice data, we examined choice run-length statistics, where runs were defined as sequences of trials in which the mouse repeated the same choice. We found that the IBL mice exhibited longer runs than if they performed the task perfectly, and the fitted GLM-HMM reproduced this feature more accurately than the classic model (Fig. ED8).

Finally, to make sure that the states identified by the model were not a consequence of the IBL mice having been previously exposed to blocks of trials with a consistent side bias (see Section 4.4), we examined data from 4 mice that were never exposed to “bias blocks”. In Fig. ED3, we show that the retrieved states, dwell times and model comparison results for these animals were similar to those shown in the rest of the IBL population (shown in Fig. 4).

### 2.5 Data provide evidence for discrete, not continuous, states

The GLM-HMM describes perceptual decision-making in terms of discrete states that persist for many trials in a row before switching. However, the model’s dramatic improvement over classic models does not guarantee that the states underlying decision-making are best described as discrete. One could imagine, for example, that a state governing the animal’s degree of engagement drifts gradually over time, and that the GLM-HMM simply divides these continuous changes into discrete clusters. To address this possibility, we fit the data with PsyTrack, a psychophysical model with continuous latent states [35, 36]. The PsyTrack model describes sensory decision-making using an identical Bernoulli GLM, but with dynamic weights that drift according to a Gaussian random walk (see Methods sec. 4.3).

For all 37 mice in our dataset, the 3-state GLM-HMM achieved substantially higher test log-likelihood than the PsyTrack model (left panel of Fig. 4f). Model selection also correctly identified simulated data from the GLM-HMM, whereas datasets simulated from a matched first-order autoregressive model had roughly equal log-likelihood under the two models (middle and right panels of Fig. 4f). Not only was the GLM-HMM favored over PsyTrack for IBL mice, but we show that it was favored over PsyTrack for the other mouse datasets we study in this work: see Fig. ED4. Finally, to verify that the GLM-HMM’s improved performance compared to PsyTrack was not merely due to the fact that the GLM-HMM does better at accounting for lapses, we also show the average fits of both PsyTrack and the 3 state GLM-HMM to the empirical choice data of the example mouse in Fig. S7. Despite the absence of explicit lapse parameters in both of these models, both are able to capture the non-zero error rates of the mouse on easy trials.

### 2.6 Mice switch between multiple strategies in a second task

To ensure that our findings were not specific to the IBL task or training protocol, we examined a second mouse dataset with a different sensory decision-making task. Odoemene *et al*. [20] trained mice to report whether the flash rate of an LED was above or below 12Hz by making a right or left nose poke (Fig. 5a). Once again, we found that the multi-state GLM-HMM provided a far more accurate description of mouse decision-making than a basic GLM or the classic lapse model (Fig. 5b). Although the performance of the 3-state and 4-state models was similar, we focused on the 4-state model because—in addition to having slightly higher test log-likelihood for a majority of animals (see Fig. S8 for individual curves)—the 4-state model balanced simplicity and interpretability, with each state in the 4-state model corresponding to a distinct behavioral strategy. (See Fig. S9 and Fig. S10 for a comparison to 3-state and 5-state fits). The 4-state model exhibited an average improvement of 0.025 bits/trial over the classic lapse model, making the test dataset approximately 1 × 10^18^ times more likely under the GLM-HMM than to the lapse model.

**Figure 5:**
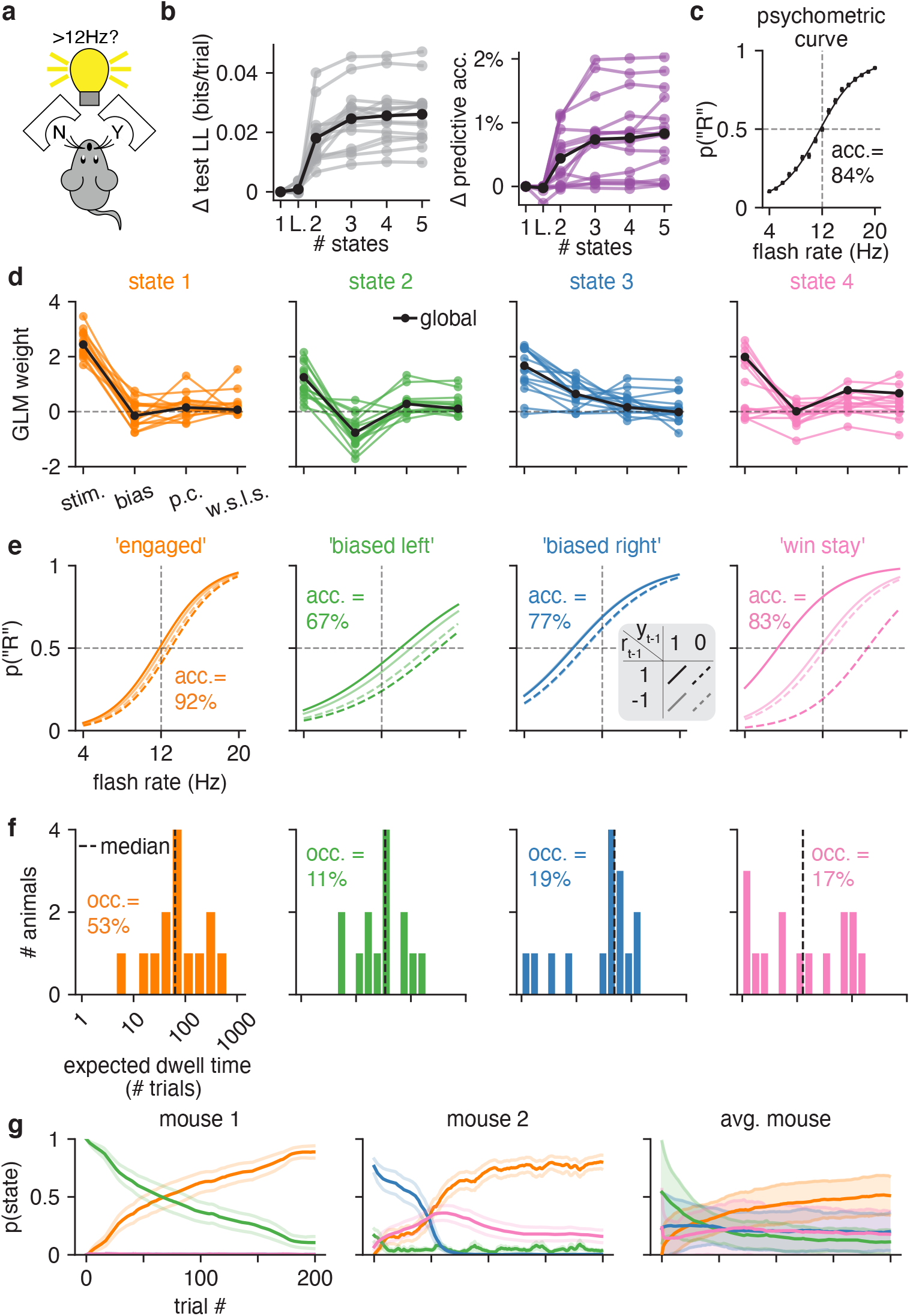
GLM-HMM application to second mouse dataset. **(a)** Mice in the Odoemene *et al*. [20] task had to indicate if the flash rate of light pulses from an LED panel was above or below 12Hz by nose poking to the right or left. **(b)** Left: test set log-likelihood for 15 mice in this dataset. Black is the mean across animals. Right: change in test set predictive accuracy relative to a GLM for each mouse in the dataset. Black is the mean across all animals. **(c)** The psychometric curve when all data from all mice are concatenated together. **(d)** The retrieved weights for a 4 state GLM-HMM for each mouse. The covariates influencing choice in each of the four states are the stimulus frequency, the mouse’s bias, its choice on the previous trial (‘p.c’) and the product of reward and choice on the previous trial (‘w.s.l.s.’). Black indicates the global fit across all mice. **(e)** The probability that the mouse went rightward in each of the four states as a function of the stimulus, previous choice and reward on the previous trial (each of the four lines corresponds to a different setting of previous choice and previous reward; the four lines are calculated using the global fit weights shown in d). We also report animals’ task accuracy (labeled ‘acc.’) when in each of the four states. As discussed in text, we label the fourth state the “win stay” state as the psychometric curve for this state is shifted so that the mouse is likely to repeat a choice only if the previous trial was rewarded.**(f)** The expected dwell times for each mouse in each state, as obtained from the inferred transition matrices (for the full transition matrices for all mice, see Fig. S11). The dashed black line indicates the median dwell time across animals. We report the fractional occupancies (labeled ‘occ.’) of the four states across all mice. For context, the median session length was 683 trials. (**g**) Left and middle: average (across 20 sessions) posterior state probabilities for the first 200 trials within a session for two example mice; error bars are standard errors. Right: posterior state probabilities for first 200 trials when averaged across all mice. Error bars indicate standard deviation across mice.

Fig. 5d shows the inferred GLM weights associated with each state in the 4-state GLM-HMM. Based on the qualitative differences in the state-specific psychometric curves (Fig. 5e), we labeled the four states as “engaged”, “biased left”, “biased right” and “win-stay”. The combination of stimulus and choice history weights for this fourth state gave rise to a large separation between psychometric curves conditioned on previous reward; the resulting strategy could be described as “win-stay” because the animal tended to repeat a choice if it was rewarded. (However, it did *not* tend to switch if a choice was unrewarded). Accuracy was highest in the engaged state (92%) and lowest in the biased-left (67%) and biased right states (77%). Similar to the IBL dataset, the identified states tended to persist for many trials in a row before switching (Fig. 5f and (Fig. ED5b)).

Finally, we used the fitted model to examine the temporal evolution of latent states within a session. Figure 5g shows the average posterior state probabilities over the first 200 trials in a session for two example mice and the average over all mice (Fig. ED6 shows average posterior state probabilities for each individual mouse in the cohort separately). These trajectories reveal that mice typically began a session in one of the two biased states, and had a low probability of entering the engaged state within the first 50 trials: mice used these initial trials of a session to “warm-up” and then gradually improved their performance [14]. This represents a substantial departure from the IBL mice, most of which had a high probability of engagement from the very first trial, and had relatively flat average trajectories over the first 90 trials of a session (Fig. 3 and Fig. S12). Note however that the warm-up effect was not the only form of state switching observed in the Odoemene *et al* mice, as the animals switched states multiple times per session and continued switching beyond the first 100 trials (Fig. ED5). We also examined whether the effects of fatigue or satiety could be observed in the average state probabilities at the end of sessions, but did not find consistent patterns across animals (Fig. ED7).

### 2.7 External correlates of engaged and disengaged states

One powerful feature of the GLM-HMM is the fact that it can be used to identify internal states from binary decision-making data alone. However, it is natural to ask whether these states manifest themselves in other observable aspects of mouse behavior. In other words, does a mouse behave differently when it is in the engaged state than in a biased state, above and beyond its increased probability of making a correct choice? To address this question, we examined response times and violation rates, two observable features that we have did not incorporate into the models.

Previous literature has revealed that humans and monkeys commit more errors on long duration trials than short duration trials [37–39]. This motivated us to examine the distributions of response times for the engaged state, compared to for the disengaged (left and right biased) states.

In the IBL dataset, we found that engaged and disengaged response time distributions were statistically different for all 37 mice (Komogorov-Smirnov tests reject the null hypothesis with p < 0.05 for all 37 mice; Fig. 6a). Examining the Q-Q plots, it was also clear that the most extreme response times for each animal were associated with the disengaged states (Fig. 6b). In the Odoemene *et al*. [20] dataset, we examined differences in the rate of violations—trials where the mouse failed to make a decision within the response period.The mean violation rate across all mice was 21% (much higher than the 1% violation rate observed in the IBL dataset), and violations were 3.2% more common in the disengaged states (2, 3, and 4) than in the engaged state (state 1).

**Figure 6:**
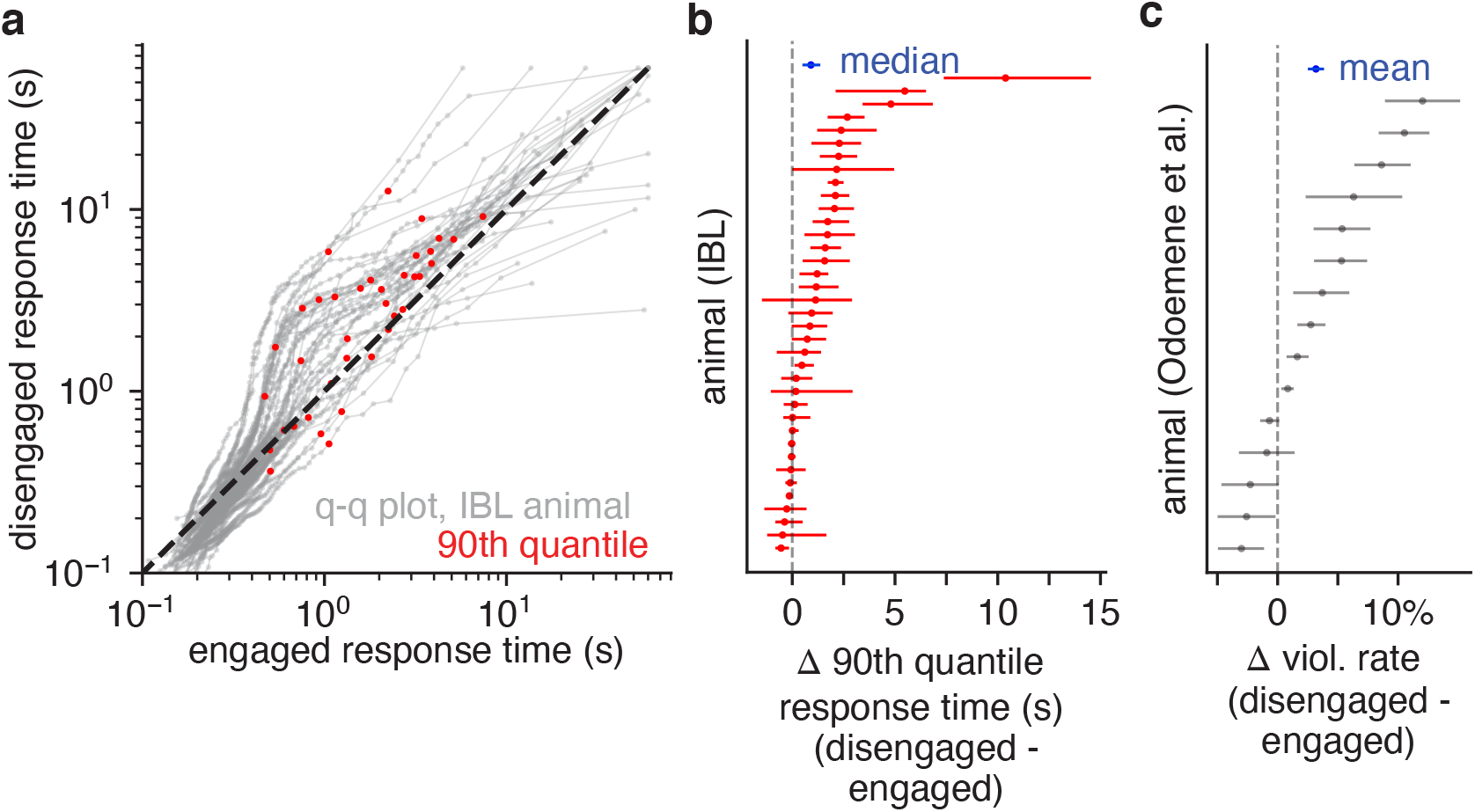
Behavioral correlates for GLM-HMM states. **(a)** Q-Q plots for response time distributions associated with engaged and disengaged (biased left/rightward) states for IBL mice. Each curve is an individual animal, and the red dots indicate the 90th quantile response times. For all 37 mice, a KS-test indicated that the engaged and disengaged response time distributions were statistically different. Furthermore, as can readily be observed from the Q-Q plots, the longest response times typically occurred in the disengaged state. **(b)** Difference in the 90th quantile response time for the engaged and disengaged states for each IBL animal, as well as 95% bootstrap confidence intervals (n=2000). Blue is the median difference across mice (0.95s), as well as the 95% bootstrap confidence interval. **(c)** Violation rate differences for mice in the Odoemene *et al*. [20] dataset when in the engaged (state 1) and disengaged states (states 2, 3, 4). Error bars are 95% bootstrap confidence intervals (n=2000); blue indicates the mean difference in violation rate across all mice (3.2%).

### 2.8 State dependent decision-making in human visual task

Finally, while the primary focus of this paper is identifying the discrete latent strategies underpinning mouse decision-making, the GLM-HMM framework is general and can be used to study choice behavior across species. We, thus, applied the GLM-HMM to the choice behavior of humans performing the visual motion discrimination task of [27]. Participants had to judge the difference in motion coherence between two consecutive random dot kinematograms: the first kinetogram always corresponded to a reference stimulus with 70% motion coherence, while the second stimulus could have greater or less motion coherence (Figure 7a). Participants indicated the stimulus they perceived to have the greater motion coherence by pressing a button. Cross-validation revealed that the choice data of 24 out of 27 humans was better explained by a 2 or 3 state GLM-HMM compared to the classic lapse model (Fig. 7b-c). The mean improvement of the 2 state GLM-HMM compared to the classic lapse model was 0.013 bits/trial, making a test dataset of 500 trials 90 times more likely to have been generated by a 2 state GLM-HMM than by the classic lapse model.

**Figure 7:**
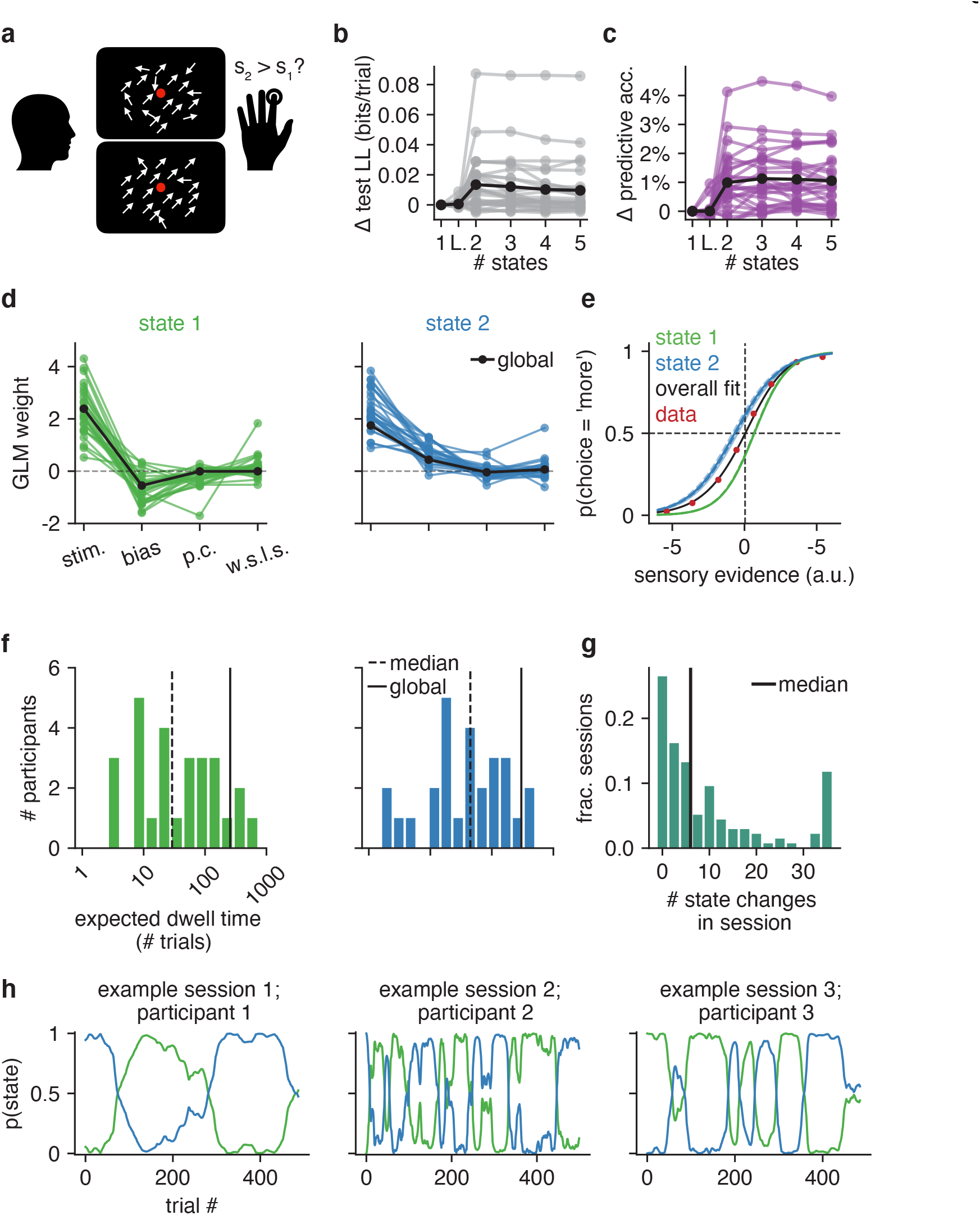
GLM-HMM application to human dataset. **(a)** 27 human participants performed the motion coherence discrimination task of [27], where they used a button to indicate if there was greater motion coherence in the first or second stimulus presented within a trial. **(b)** Change in test set log-likelihood relative to a GLM for each of the 27 participants. Black is the mean across participants. **(c)** Change in test set predictive accuracy relative to a GLM for each participant in the dataset. Black is the mean across participants. **(d)** Retrieved weights for a 2 state GLM-HMM for each participant. The covariates influencing choice in each of the four states are the difference in motion coherence between the two stimuli, the participant’s bias, their choice on the previous trial (‘p.c’) and the product of reward and choice on the previous trial (‘w.s.l.s.’). Black indicates the global fit across all participants. **(e)** Similar to Figs. 2g and 5e, the blue and green curves give the probability that the participant went rightward in each of the two states as a function of the stimulus, previous choice and previous reward. That all blue curves are overlapping indicates the lack of dependence on previous choice and reward in this state; similarly for state 1. The black curve is the overall psychometric curve generated by the GLM-HMM. Specifically, using the global GLM-HMM parameters, we generated the same number of choices as those in the human dataset. At each trial, regardless of the true stimulus presented, we calculated *p*_*t*_(“*R*”) for each possible stimuli by averaging the per-state psychometric curves (blue and green) and weighting by the appropriate row in the transition matrix (depending on the sampled latent state at the previous trial). Finally, we averaged the per-trial psychometric curves across all trials to obtain the curve that is shown in black, while the red dots represent the empirical choice data. **(f)** The expected dwell times for each participant in each state, as obtained from the inferred transition matrices. The black dashed line indicates the median dwell time across participants, while the black solid line indicates the global fit (see Algorithm 1). **(g)** Empirical number of state changes per session obtained using posterior state probabilities such as those shown in h; median session length is 500 trials. Black indicates the median number of state changes across all sessions. **(h)** Posterior state probabilities for three example sessions corresponding to three different participants.

Fig. 7d shows the retrieved weights for each participant for the 2 state model (we focus on the 2 state model as this was the model favored by the majority of participants), while Fig. 7e shows the probability of the participant selecting, in each of the two states (blue and green curves), that the second stimulus had greater motion coherence than the first (choice = ‘more’) as a function of the relative motion coherence. The 2 states correspond to participants having a bias for selecting each of the two possible outcomes - when using the blue state, participants were biased toward selecting ‘more’, while participants were biased toward selecting ‘less’ in the green state. Fig. 7f shows the expected dwell times for all participants in each state, while Fig. 7g shows the empirical number of state changes detected in each session (across the entire participant population). The median number of state changes per session was 6 switches, indicating that it is not just mice that switch between strategies multiple times per session. Fig. 7h illustrates the strategy changes across a session and shows the posterior state probabilities for 3 example sessions (for 3 different participants).

## 3 Discussion

In this work, we used the GLM-HMM framework to identify hidden states from perceptual decision-making data. Mouse behavior in two different perceptual decision-making tasks [19, 20] was far better described by a GLM-HMM with sustained engaged and biased states. Unlike the classic lapse model, these states alternated on the timescale of tens to hundreds of trials. We also applied the model to human psychophysical data [27] and found that human behavior was also better described by a GLM-HMM with sustained states that differed in bias. While it may seem obvious that mice should exhibit fluctuations in task engagement over time, this is a very different account of behavior to that given by the classic lapse model, which assumes that error trials do not occur in blocks, and are instead interspersed throughout a session. Furthermore, our method offers the practical benefit that it allows practitioners to segment trials according to the inferred behavioral strategy. Finally, it was not obvious that this alteration in choice policy over the course of a session should occur in a discrete rather than continuous manner. Comparing our results with those for PsyTrack—a model with continuous decision-making states— provided support for the view that mice use discrete rather than continuous decision-making states.

While we found similarities in the strategies pursued by mice performing the different visual detection tasks, we also found some differences. For example, we found that mice performing the Odoemene *et al*. [20] task used an additional ‘win-stay’ strategy that was not observed in the IBL mice. One possible explanation for these strategy differences is that they arise due to the different shaping protocols associated with the different tasks. A follow-up study could investigate the role of shaping on GLM-HMM behavioral strategies.

The ability to infer the state or strategy employed by an animal on a trial-by-trial basis will be useful for characterizing differences in performance across sessions and across animals. It will also provide a powerful tool for neuroscientists studying the neural mechanisms that support decision-making [40–46]. It may be the case that different strategies rely on different circuits or different patterns of neural activity [47].

Although we found evidence of warm-up behavior at the start of sessions in the Odoemene *et al*. [20] mice, we were somewhat surprised to find few signatures of satiation or fatigue toward the end of a session. This might be due to the fact that sessions were typically of fixed duration, and may have ended before mice had a chance to grow satiated or fatigued. One future direction will be to apply the GLM-HMM to experiments with longer sessions, where it may be useful for detecting changes in behavior reflecting satiety or fatigue.

Compared to the mice we studied, humans used two relatively engaged strategies with opposite biases when performing the perceptual decision-making task of [27]. Unfortunately, it is impossible to say if the difference in strategies observed across species were due to species differences, or to differences in task protocol. An interesting follow-up experiment would be to apply the GLM-HMM to data from different species performing the same task.

In future work, we will aim to make explicit the connection between the ‘engaged’ and ‘disengaged’ strategies identified by our model and existing measures of arousal and engagement in the literature. Identifying the relationship between the GLM-HMM’s hidden states and pupil diameter, low-frequency LFP oscillations, spontaneous neuronal firing, noise correlations and the action of neuromodulators [48–51] will be a priority.

That discrete states underpin mouse and human choice behavior may also call for new normative models to explain why subjects develop these states to begin with [52]. The existence of disengaged and engaged states could reflect explore-exploit behavior [17, 53, 54], optimal learning (e.g., [55–58]), or could simply indicate incomplete learning of the task. Alternatively, if the states uncovered by our model reflect a similar phenomenon to the internal and external modes of sensory processing observed in [59, 60], it may be that they arise due to the limited information-processing capacity of the brain.

Another promising direction will be to replace the model’s fixed transition matrix with a generalized linear model so as to allow external covariates to modulate the probability of state changes[6]. This would allow us to identify the factors that influence state changes (e.g., a preponderance of unrewarded choices) and, potentially, seek to control such transitions.

While there are many avenues for future research, we believe that these results call for a significant rethinking of rodent and human perceptual decision-making behavior and the methods for analyzing it. Indeed, standard analysis methods do not take account of the possibility that an animal makes abrupt changes in decision-making strategy multiple times per session. We feel that the ability to infer internal states from choice behavior will open up new directions for data analysis and provide new insights into a previously inaccessible dimension of perceptual decision-making behavior.

## 4 Methods

We confirm that our research complies with all relevant ethical regulations.

### 4.1 Inference of GLM-HMM parameters

#### 4.1.1 GLM-HMM objective function

When fitting the GLM-HMM, the goal is to learn the transition matrix *A* ∈ ℝ ^*K*×*K*^, the set of weights influencing choice in each state,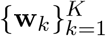 with **w**_*k*_ ∈ ℝ ^*M*^, and the initial state distribution, ***π*** ∈ ℝ ^*K*^. We fit this set of parameters, collectively labeled as Θ, to choice data using Maximum A Posteriori (MAP) estimation. This was practically implemented by the Expectation Maximization (EM) algorithm [61].

The EM algorithm has been adapted to fit Hidden Markov models with external inputs before [6, 31, 32]. However, there are some application-specific choices in the implementation, so for completeness we describe the full procedure here.

The EM algorithm seeks to maximize the log-posterior of the parameters given the choice and input data, 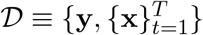. The log-posterior is given, up to an unknown constant, by:

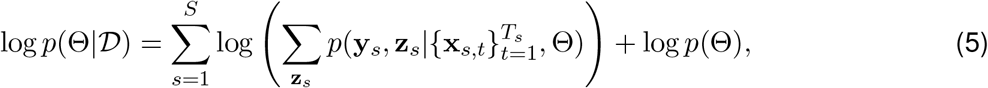

where *s* indexes the session in which the data was collected (out of *S* total sessions), and the sum over **z**_*s*_ is over all 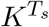 possible latent state paths in the *T*_*s*_ trials of session *s*.

The prior distribution over the model parameters *p*(Θ) that we used was:

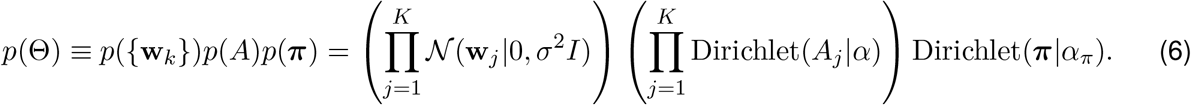

The prior over the GLM weight vectors **w**_*k*_ was thus an independent, zero-mean Gaussian with variance *σ*^2^. Smaller values of *σ*^2^ have the effect of shrinking the fitted weights towards zero, while *σ*^2^ = ∞ corresponds to a flat prior. For the transition matrix, we placed an independent Dirichlet prior over each row *A*_*i*_, which is a natural choice for vectors on the unit (*K* − 1) simplex (i.e., all elements of the vector must be non-negative and sum to 1). The Dirichlet distribution is controlled by a shape parameter 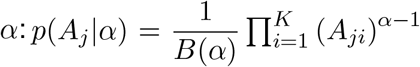. A value of *α* = 1 corresponds to a flat prior, while large values of *α* have the effect of spreading probability mass equally over states. We also placed a Dirichlet prior over the initial state distribution ***π***.

In order to select the hyperparameters *σ* and *α* governing the prior, we performed a grid search for *σ* ∈ {0.5, 0.75, 1, 2, 3} and *α* ∈ {1, 2} and selected the hyperparameters that resulted in the best performance on a held-out validation set. For IBL mice, the prior hyperparameters selected were *σ* = 2 and *α* = 2. For mice in the Odoemene *et al*. [20] dataset, the best hyperparameters were *σ* = 0.75 and *α* = 2. Finally, for the initial state distribution, we set *α*_*π*_ = 1.

#### 4.1.2 Expectation Maximization Algorithm

We used the Expectation Maximization (EM) algorithm [61, 62] to maximize the log posterior given in Eq. 5 with respect to the GLM-HMM parameters. As the sum given in Eq. 5 involves an exponential number of terms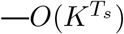 terms to be specific—we don’t maximize this expression directly. Instead, the EM algorithm provides an efficient way to compute this term using a single forward and backward pass over the data. During the E-step of the EM algorithm we compute the “Expected Complete-Data log-likelihood” (ECLL), which is a lower bound on the right-hand side of Eq. 5 [61–63]. Then, during the ‘Maximization’ or M-step of the algorithm, we maximize the ECLL with respect to the model parameters, Θ. It can be shown that this procedure has the effect of always improving the log posterior in each step of the algorithm [63].

Concretely, the ECLL can be written as a sum over the ECLLs for each session and the log-prior. The ECLL for a single session *s* can be written

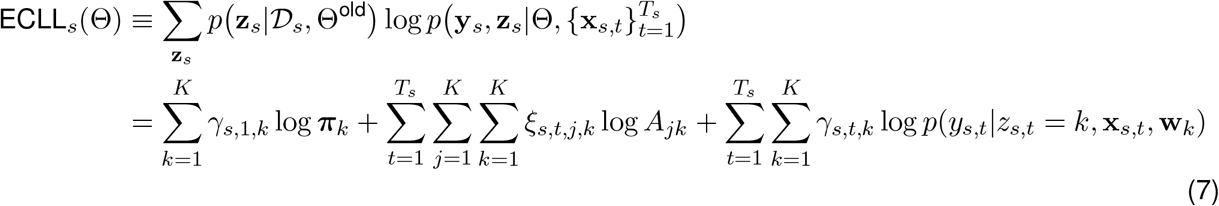

In order to get to the second line, we substituted the definition of the joint distribution for the GLM-HMM:

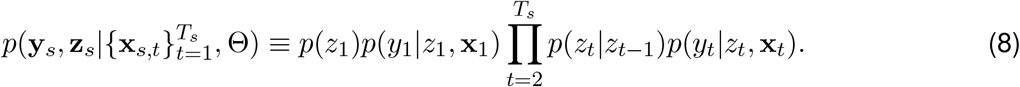

In Eq. 7, *p*(*y*_*s,t*_|*z*_*s,t*_ = *k*, **x**_*s,t*_, **w**_*k*_) is the Bernoulli GLM distribution given by Eq. 4. Finally,

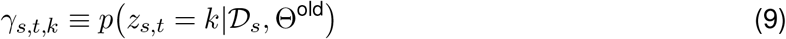

is the posterior state probability at trial *t* (in session *s*) for state *k*, while

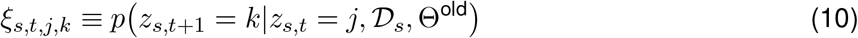

is the joint posterior state distribution for two consecutive latents.

While the formula for the log posterior (Eq. 5) involved the sum over all possible state assignments at each trial, by taking advantage of the structure of the joint probability distribution for the GLM-HMM (Eq. 8), this sum was implemented efficiently and Eq. 7 involves summing over at most *O*(*K*^2^*T*_*s*_) elements.

The single and joint posterior state probabilities, *γ*_*s,t,k*_ and *ξ*_*s,t,k*_, are estimated via the forward-backward algorithm, which we describe below.

#### 4.1.3 Expectation Step: Forward-Backward Algorithm

During the E-Step of the EM algorithm, the single and joint posterior state probabilities for all trials and states, {*γ*_*s,t,k*_} and {*ξ*_*s,t,j,k*_}, are estimated using the forward-backward algorithm [64] at the current setting of the GLM-HMM parameters, Θ^old^.

The forward-backward algorithm makes use of recursion and memoization to allow these posterior probabilities to be calculated efficiently, with the forward and backward passes of the algorithm each requiring just a single pass through all trials within a session.

The goal of the forward pass is to obtain, for each trial *t* within session *s* and each state *k*, the quantity

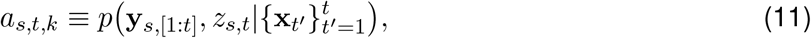

which represents the posterior probability of the choice data up until trial *t* and the latent state at trial *t* being state *k*.

The posterior probability associated with trial 1, *a*_*s*,1,*k*_, can be calculated as follows:

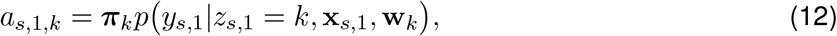

where *p* (*y*_*s*,1_|*z*_*s*,1_ = *k*, **x**_*s*,1_, **w**_*k*_) is the usual Bernoulli GLM distribution.

For trials 1 < *t* ≤ *T*_*s*_, we can obtain these probabilities as follows:

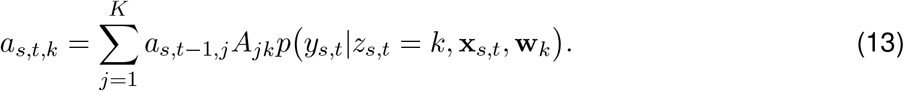

During the backward pass, the goal is to calculate the posterior probability of the choice data beyond the current trial, *b*_*s,t,k*_, for each trial *t* within session *s* and for all states, *k*:

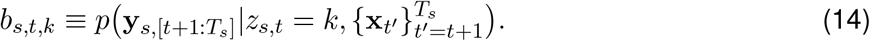

Similar to the forward pass, these quantities can be calculated efficiently by recognizing that

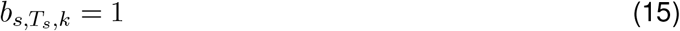

and, for *t* ∈ {*T*_*s*_ − 1, …, 1}:

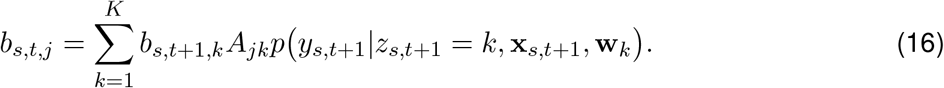

From the *a*_*s,t,k*_ and *b*_*s,t,k*_ quantities obtained via the forward-backward algorithm, we can form the single and joint posterior state probabilities, *γ*_*s,t,k*_ and *ξ*_*s,t,j,k*_, that we actually care about for forming the ECLL of Eq. 7 as follows:

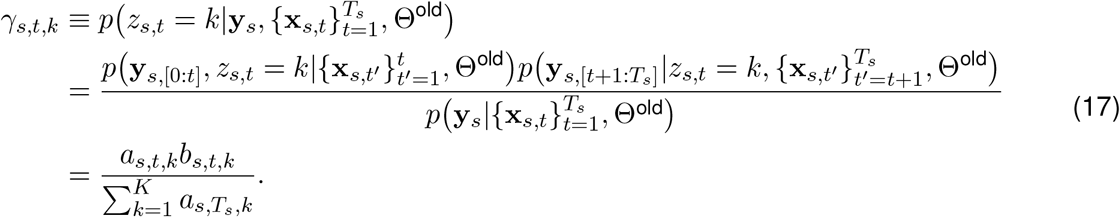

Similarly,

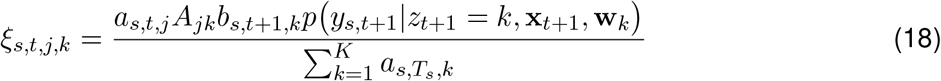

where, once again, *p*(*y*_*s,t*+1_|*z*_*t*+1_ = *k*, **x**_*t*+1_, **w**_*k*_) is the Bernoulli GLM distribution.

#### 4.1.4 Maximization Step

After running the forward-backward algorithm, we can compute the total ECLL by summing over the per-session ECLLs (Eq. 7) and adding the log prior. During the M-step, we maximize the ECLL with respect to the GLM-HMM parameters, Θ. For the initial state distribution ***π*** and the transition matrix *A*, this results in the closed-form updates:

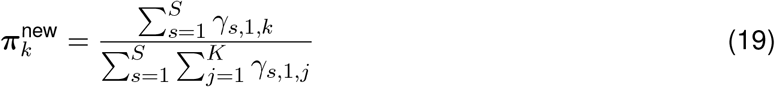

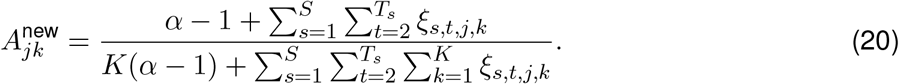

For the GLM weights, there is no such closed form update, but a Bernoulli GLM falls into the class of functions mapping external inputs to HMM emission probabilities considered in [31], so we know that the ECLL is concave in the GLM weights. As such, we can numerically find the GLM weights that maximize (not just locally but globally) the ECLL using the BFGS algorithm [65–68] as implemented by the scipy optimize function in python [69].

#### 4.1.5 Comparing states across animals and GLM-HMM parameter initialization

In Fig. 4 and Fig. 5, we show the results from fitting a single GLM-HMM to each animal; however it is nontrivial to map the retrieved states across animals to one another. As such, we employed a multistage fitting procedure that allowed us to make this comparison, and we detail this procedure in Algorithm 1. In the first stage, we concatenated the data for all animals in a single dataset together (for the IBL dataset, this would be the data for all 37 animals). We then fit a GLM (a 1 state GLM-HMM) to the concatenated data using Maximum Likelihood estimation. We used the fit GLM weights to initialize the GLM weights of a *K*-state GLM-HMM that we again fit to the concatenated dataset from all animals together (to obtain a “global fit”). We added Gaussian noise with *σ*_init_ = 0.2 to the GLM weights, so that the initialized states were distinct, and we initialized the transition matrix of the *K*-State GLM-HMM as 0.95 × 𝕝+ 𝒩 (0, Σ_trans._) where 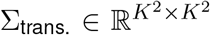 and Σ_trans._ = 0.05 × 𝕝. We then normalized this so that that rows of the transition matrix added up to 1, and represented probabilities. While the EM algorithm is guaranteed to converge to a local optimum in the log probability landscape of Eq. 5, there is no guarantee that it will converge to the global optimum [70]. Correspondingly, for each value of *K*, we fit the model 20 times using 20 different initializations.

In the next stage of the fitting procedure, we wanted to obtain a separate GLM-HMM fit for each animal, so we initialized a model for each animal with the GLM-HMM global fit parameters from all animals together (out of the 20 initializations, we chose the model that resulted in the best training set log-likelihood). We then ran the EM algorithm to convergence; it is these recovered parameters that are shown in Fig. 4 and Fig. 5. By initializing each individual animal’s model with the parameters from the fit to all animals together, it was no longer necessary for us to permute the retrieved states from each animal so as to map semantically similar states to one another.

##### Algorithm 1 Multistage GLM-HMM fitting procedure

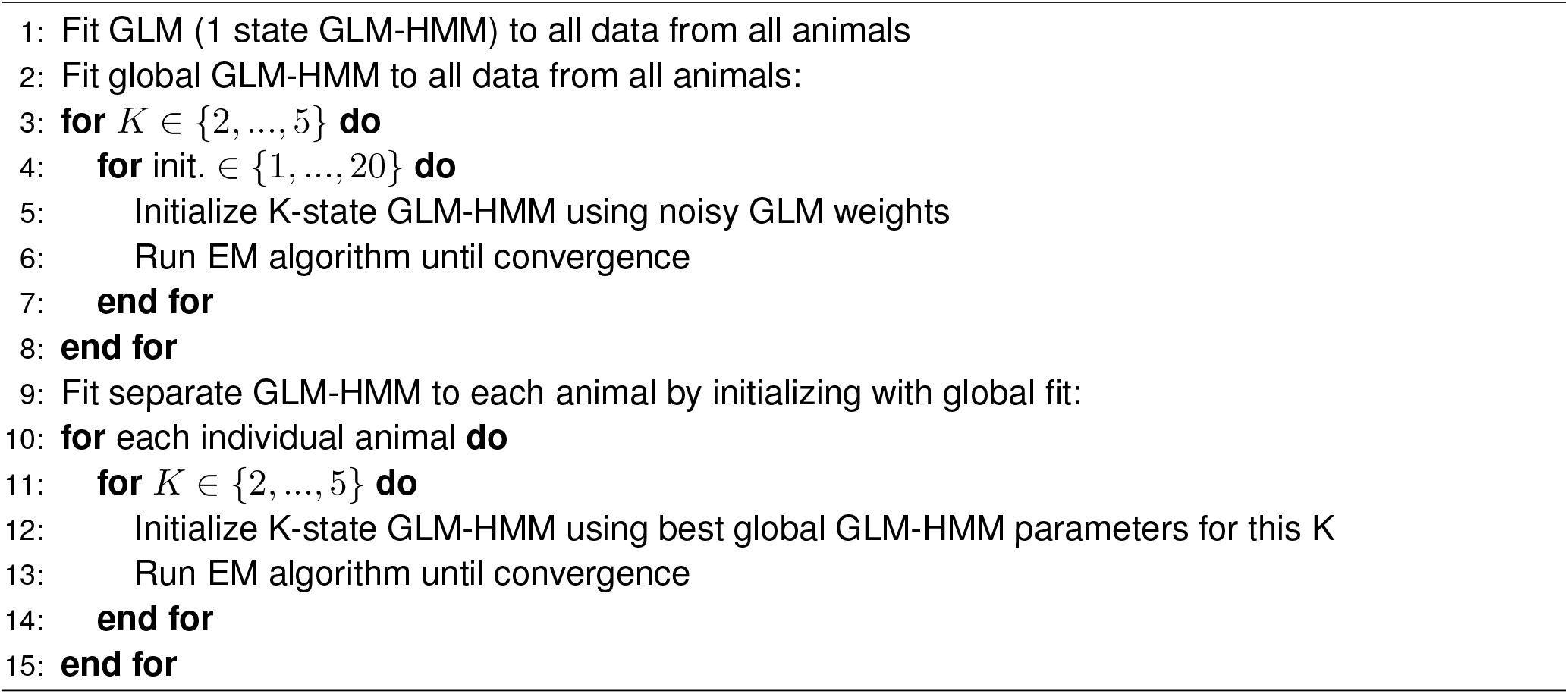

We note that the initialization scheme detailed above is sufficiently robust so as to allow recovery of GLM-HMM parameters in various parameter regimes of interest. In particular, we simulated datasets from a GLM-HMM with the global fit parameters for both the IBL and Odoemene *et al*. [20] datasets, as well as a global fit lapse model. We show the results of these recovery analyses in Fig. ED9 and Fig. ED10.

### 4.2 Assessing Model Performance

#### 4.2.1 Cross Validation

There are two ways in which to perform cross-validation when working with Hidden Markov Models. Firstly, it is possible to hold out entire sessions of choice data for assessing test-set performance. That is, when fitting the model, the objective function in Eq. 5 and the ECLL in Eq. 7 are modified to only include 80% of sessions (since we use 5-fold cross-validation throughout this work); and the log-likelihood of the held-out 20% of sessions is calculated using the fit parameters, and a single run of the forward-pass on the held-out sessions:

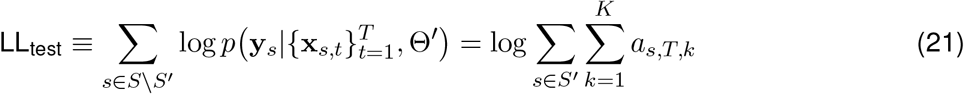

where *S* \ *S*′ is the set of held out sessions, and Θ′ is the set of GLM-HMM parameters obtained by fitting the model using the trials from *S*′.

The second method of performing cross-validation involves holding out 20% of trials within a session. When fitting the model, the third term in the ECLL is modified so as to exclude these trials and is now 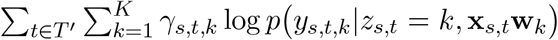, where *T*′ is the set of trials to be used to fit the model. Furthermore, the calculation of the posterior state probabilities, *γ*_*s,t,k*_ and *ξ*_*s,t,j,k*_, is also modified so as to exclude the test set choice data. In particular, *γ*_*s,t,k*_ is now *p*(*z*_*s,t*_|{*y*_*s,t*′_}_*t*′ ∈ *T*′_, {**x**_*s,t*′_}_*t*′ ∈*T*′_, Θ^old^) and similarly *ξ*_*s,t,j,k*_ is now *p* (*z*_*s,t*+1_ = *k*|*z*_*s,t*_ = *j*, {*y*_*s,t*′_}_*t*′ ∈*T*′_, {**x**_*s,t*′_}_*t*′ ∈*T*′_, Θ^old^). The method of calculating these modified posterior probabilities is as detailed in Eq. 17 and Eq. 18, but now the calculation of the forward and backward probabilities, *a*_*s,t,k*_ and *b*_*s,t,k*_ in Eq. 12, Eq. 13, Eq. 15 and Eq. 16 is modified so that, on trials that are identified as test trials, the *p* (*y*_*s,t*_|*z*_*s,t*_ = *k*, **x**_*s,t*_, **w**_*k*_) term in these equations is replaced with 1.

In Fig. 2, Fig. 4 and Fig. 5, we perform cross-validation by holding out entire sessions. We believed it would be harder to make good predictions on entire held out sessions, compared to single trials within a session, as we thought that mice would exhibit more variability in behavior across sessions compared to within sessions. When we compare the performance of the GLM-HMM against the PsyTrack model of [35] in Fig. 4f, we use the second method of cross-validation so as to use the same train and test sets as PsyTrack (PsyTrack cannot make predictions on entire held-out sessions).

#### 4.2.2 Units for test set log-likelihood

In Fig. 2, Fig. 4, Fig. 5 and Fig. 7, as well as in some supplementary figures, we report the log-likelihood of different models on held-out sessions in units of bits per trial. This is calculated as follows:

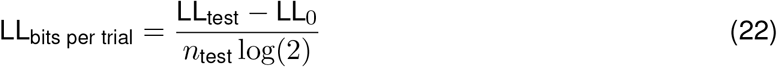

where, for the GLM-HMM, LL_test_ is the log-likelihood of the entire test set as calculated in Eq. 21, and LL_0_ is the log-likelihood of the same test set under a Bernoulli model of animal choice behavior. Specifically, this baseline model assumes that animals flip a coin on each trial so as to decide to go Right, and the probability of going Right is equal to the fraction of trials in the training set in which the animal chose to go to the Right. *n*_test_ is the number of trials in the test set, and is important to include since LL_test_ depends on the number of trials in the test set. Dividing by log(2) gives the log-likelihood the units of bits per trial. Clearly, larger values of the log-likelihood are better, with a value of 0 indicating that a model offers no improvement in prediction compared to the crude baseline model described above. However, even small values of test set log-likelihood, when reported in units of bits per trial, can indicate a large improvement in predictive power. For a test set size of *n*_test_ = 500, a log-likelihood value of 0.01 bits/trial indicates that the test data is 31.5 times more likely to have been generated with the GLM-HMM compared to the baseline model. For a test set of *n*_test_ = 5000, and the same value of 0.01 bits/trial for the log-likelihood, the test set becomes 1 × 10^15^ times more likely under the GLM-HMM compared to the baseline model!

#### 4.2.3 Predictive Accuracy

In Fig. 2, Fig. 4 and Fig. 5, we also report the predictive accuracy of the GLM-HMM. When calculating the predictive accuracy, we employ a method similar to the second method described above in section 4.2.1. In particular, we hold out 20% of trials and then obtain the posterior state probabilities for these trials, *t*″ ∈ {*T* \ *T*′}, as *γ*_*s,t*″, *k*_ = *p* (*z*_*s,t*″_ | {*y*_*s,t*′_} _*t*′ ∈*T*′_, {**x**_*s,t*′_}_*t*′ ∈*T*′_, Θ), using the other 80% of trials (this latter set of trials being labeled *T*′). We then calculate the probability of the held-out choices being to go Right as:

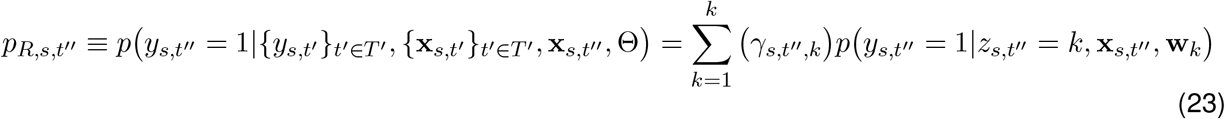

We then calculate the predictive accuracy as:

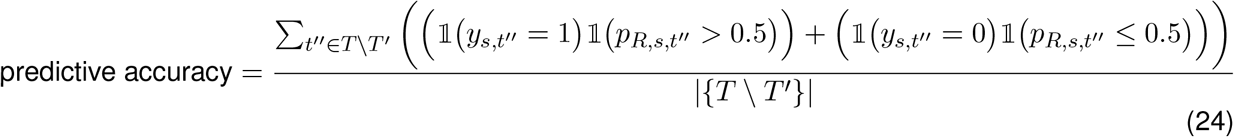

### 4.3 Comparison with PsyTrack Model of Roy et al

The PsyTrack model of Roy et al. [36] assumes an animal makes its choice at trial *t* according to

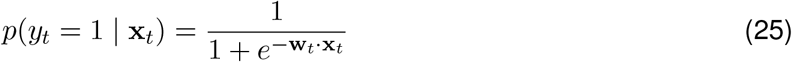

where **w**_*t*_ evolves according to

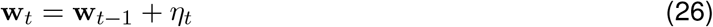

where 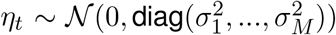 and 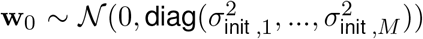. Specifically, the animal is assumed to use a set of slowly changing weights to make its decision on each trial.

In order to perform model comparison with the PsyTrack model of [35, 36], we utilized the code provided at https://github.com/nicholas-roy/psytrack (version 1.3).

#### 4.3.1 Simulating Choice Data with AR(1) Model for Weights

While the GLM-HMM is a generative model that can be readily used to simulate choice data that resembles, in accuracy and in the resulting psychometric curves, the choice data of real animals, this is not true for the PsyTrack model of [35, 36]. Indeed, specifying only the 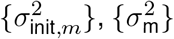 hyperparameters of that model and then generating weights according to Eq. 26 and choice data according to Eq. 25 will likely result in choice behavior that is vastly different from that of real animals (the PsyTrack model is underconstrained as a generative model). As such, so as to produce the rightmost panel of Fig. 4f, we simulated smoothly evolving weights from an AR(1) model where the parameters of this model were obtained using the PsyTrack fits to real data. Specifically, we assumed that the probability of going rightward at time t was given by:

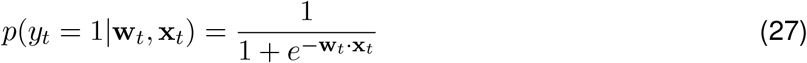

and we assumed that the weights evolved according to an AR(1) process as follows:

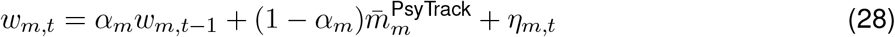

where *w*_*m,t*_ is the *m*th element of **w**_*t*_ in Eq. 27 and 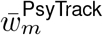 is the average weight that an animal places on covariate *m* across all trials when fit with the PsyTrack model: 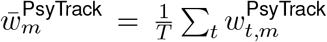. We obtained *α*_*m*_ by regressing 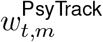 against 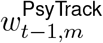 and taking the retrieved slope (after confirming that the retrieved slope had a magnitude of less than 1, so as to ensure that the weights did not diverge as *t* → ∞).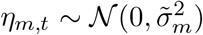, where 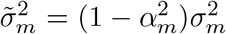 and 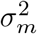 was obtained from the PsyTrack fit to an animal’s real choice data. Finally, we set **w**_0_ = 0.

We simulated weight trajectories for thousands of trials for each animal, so that the AR(1) process reached the stationary regime for each covariate, and so that the mean, variance and autocovariance of each weight for each covariate were close to those returned by the PsyTrack fits to the real choice data.

### 4.4 Datasets studied

In this paper, we applied the GLM-HMM to three publicly available behavioral datasets associated with recent publications. Firstly, we studied the data associated with [19] that is made available via figshare at https://doi.org/10.6084/m9.figshare.11636748. We used the framework developed in [71] to access the data. We modeled the choice data for the 37 animals in this dataset which had more than 30 sessions of data during the ‘bias block’ regime. We focused on this regime because of the fact that mice, when they have reached this regime, understand the rules of the task and exhibit stationary behavior (see Fig. S1 for plots of accuracy against session identity for each animal, as well as Fig. S2 for the psychometric curves for these animals for the trials studied). For each session, we subset to the first 90 trials of data because, during these trials, the stimulus was equally likely to appear on the left or right of the screen. After the first 90 trials, the structure of the task changed and for a block of trials, the stimulus appeared on the left with a probability of either 80% or 20%; the block identity switched multiple times throughout a session, so that 80% and 20% blocks were interleaved. We subset to the animals with more than 30 sessions of data because we were able to confidently recover GLM-HMM and lapse model generative parameters when we simulated datasets with this number of trials: see Figures Fig. ED9 and Fig. ED10. As a sanity check to make sure that the recovered states and transitions were not a consequence of the animals we study having been exposed to bias blocks in earlier sessions, we obtained data for 4 animals that were never exposed to bias blocks (not included in the publicly released dataset) and fit the GLM-HMM to the choice data for these animals. In Fig. ED3, we show that the retrieved states, dwell times and model comparison results for these animals look very similar to those shown in Fig. 4.

The second dataset that we studied was that associated with [20], with the data being made available at https://doi.org/10.14224/1.38944. Once again, we studied sessions after animals had learned the task (see Fig. S13 and Fig. S14). For this dataset, the retrieved states were less distinct compared to those for the IBL dataset, and as such, we required more trials to be able to recover the generative parameters in simulated data: see Fig. ED9. We thus subset to the 15 animals with more than 20 sessions of data and 12,000 trials of data. Compared to the IBL dataset, where the violation rate across all animals’ data was less than 1% of trials (where a violation is where the animal chose not to respond), the violation rate across the 15 animals that we studied from this second dataset was 21%. Thus, it was important to develop a principled method for dealing with violation trials. We treated violation trials as trials with missing choice data, and we handled these trials in a similar way to how we handled test data when performing the second type of cross-validation described in section 4.2.1 above. That is, we modified the third term of the ECLL given in Eq. 7 to exclude violation trials, and we modified the definition of the posterior state probabilities for these trials to be *γ*_*s,t,k*_ = *p* (*z*_*s,t*_|{*y*_*s,t*′_}_*t*′ ∈*T*′_, {**x**_*s,t*′_}_*t*′ ∈*T*′_, Θ^old^) and *ξ*_*s,t,j,k*_ = *p*(*z*_*s,t+1*_=*k*|*z*_*s,t*_ = *j*, {*y*_*s,t*′_}_*t*′ ∈ *T*′_, {**x**_*s,t*′_}_*t*′ ∈ *T*′_, Θ^old^), where *T* ′, rather than representing the training set data, is now the set of non-violation trials. The calculation of the forward and backward probabilities, *a*_*s,t,k*_ and *b*_*s,t,k*_, was modified so that, on violation trials, in Eq. 12, Eq. 13, Eq. 15 and Eq. 16, the *p* (*y*_*s,t*_|*z*_*s,t*_ = *k*, **x**_*s,t*_, **w**_*k*_) term was replaced with 1.

Finally, the third dataset that we studied was the human dataset associated with [27], with the data being made available at https://doi.org/10.6084/m9.figshare.4300043. We applied the GLM-HMM to the data from all 27 participants in this dataset. This dataset has 5 sessions, each of approximately 500 trials, for each participant and does not include the initial session that was used to teach participants how to perform this task.

#### 4.4.1 Forming the Design Matrix

Each of the models discussed in this paper (GLM-HMM, the classic lapse model, the PsyTrack model of Roy et al. [36]) were fit using a design matrix of covariates, *X* ∈ ℝ^*T* ×*M*^, where T was the number of trials of choice data for a particular animal. A single row in this matrix was the vector of covariates, **x**_*t*_ ∈ ℝ^*M*^, influencing the animal’s choice at trial *t*. For all analyses presented in text, unless specified otherwise, *M* = 4.

For both tasks, the first column in the design matrix was the z-scored stimulus intensity. For the IBL task, we calculated the stimulus intensity as the difference in the value of the visual contrast on the right side of the screen minus the visual contrast on the left of the screen. This resulted in 9 different values for the ‘signed contrast’: {−100, −25, −12.5, −6.25, 0, 6.25, 12.5, 25, 100}. We then z-scored this difference quantity across all trials. For the Odoemene *et al*. [20] task, we subtracted the 12Hz threshold from the flash rate presented on each trial and then z-scored the resulting quantity. For the human dataset [27], we z-scored the relative motionstrength presented on each trial.

For all trials, all animals and both tasks, the second column of the design matrix was set to 1, so as to enable us to capture the animal’s innate bias for going rightward or leftward. The third column in the design matrix was the animal’s choice on the previous trial 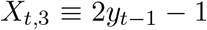. Whereas *y*_*t*−1_ ∈ {0, 1}, *X*_*t*,3_ ∈ {−1, 1}. It is not strictly necessary to perform this scaling, but we did so to ensure that the range of values for *X*_:,1_ and *X*_:,3_ were more similar (which can be useful when performing parameter optimization). Finally, the fourth column in the design matrix was the win-stay-lose-switch covariate, which was calculated as 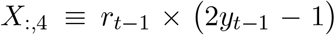, where *r*_*t*−1_ ∈ {−1, 1} was a binary variable indicating whether or not the animal was rewarded on the previous trial. Again *X*_:,4_ ∈ {−1, 1}.

#### 4.4.2 Response time data

In Fig. 6, we analyzed response times for IBL animals, as reaction time data was unavailable for most sessions. The mouse’s response time is the time from the stimulus onset to the time at which it received feedback on its decision (receiving its reward/hearing an auditory cue to indicate an error trial). In comparison, a reaction time for the IBL task is the time from the stimulus appearing on the screen to the mouse moving the wheel for the first time in the direction of its decision. Response times ranged from a few hundred milliseconds to 10 seconds in the most extreme case, while typical reaction times (calculated on a few sessions for which we had reliable data) tended to be on the order of a few hundred milliseconds. Relative to other tasks, the IBL task is a task that results in short reaction times.

#### 4.4.3 Data Availability

The raw data studied in this paper is publicly available: the IBL data associated with [19] can be accessed at https://doi.org/10.6084/m9.figshare.11636748. The Odoemene et al. data associated with [20] can be accessed at https://doi.org/10.14224/1.38944. Finally, the human data associated with [27] can be accessed at https://doi.org/10.6084/m9.figshare.4300043.

### 4.5 Code availability

We contributed code to version 0.0.1 of the Bayesian State Space Modeling framework of [72] and we use this code base to perform GLM-HMM inference. The code to analyze the resulting model fits and to produce the figures in this paper is available at https://github.com/zashwood/glm-hmm.

### 4.6 Statistics and Reproducibility

We examined the choice data of animals in the IBL dataset with more than 3000 trials of data. In the case of Odoemene et al. animals, we required 12,000 trials of data. As detailed in the Methods section, as well as in the supplementary information, we determined that these numbers were sufficient because we found that we could successfully recover the generative parameters of a GLM-HMM in simulated data in the two different parameter regimes with these numbers of trials. All animals with the requisite number of trials of data were included in the analyses we present in the paper. Each experiment presented in the paper was repeated in multiple animals (37 in the case of the IBL dataset, 15 for the Odoemene et al. dataset and 27 in the case of the Urai et al. human dataset). The effects identified were largely consistent across subjects: the choice data for all 37 IBL animals, 15 Odoemene et al. mice and for 24 out of the 27 humans were better explained by a GLM-HMM compared to a classic lapse model. Analysis was performed with code that is freely available to promote replication. We randomized the allocation of sessions to folds for each animal when performing 5-fold cross-validation. Blinding was not performed. All trials from animals with the requisite number of trials were analyzed; blinding was not necessary as cross-validation ensured that the results were not due to the particular choice of train/test split.

## 5 Acknowledgments

We are grateful to Miles Wells, Rebecca Terry, Laura Funnell and the Cortexlab at University College London for providing us with the data for the 4 mice plotted in Fig. ED3. We are grateful to Scott Linderman for developing the beautiful Bayesian State Space Modeling framework of [72]: as described in our Methods section, we built our code on top of this framework. We thank members of the Pillow Lab, the International Brain Laboratory (IBL), and specifically the Behavior Analysis Working Group within the IBL for helpful feedback throughout the project. We thank Peter Dayan, Sebastian Bruijns and Liam Paninski for acting as the IBL Review Board for this paper. We thank Emily Dennis for feedback at various points during the project. We thank Abigail Russo and Matthew Whiteway for providing feedback on drafts of this manuscript. We thank Hannah Bayer for her help and advice as we were preparing to submit this paper. Finally, we thank the anonymous reviewers for their insightful comments; our manuscript is greatly improved as a result of your input.

This work was supported by grants SCGB AWD543027 (J.W.P.) and SCGB AWD543011 (A.P.) from the Simons Collaboration on the Global Brain, grants NS104899 (J.W.P.), R01EB026946 (J.W.P.) and R01EY022979 (A.K.C.) from the NIH BRAIN initiative, U19 NIH-NINDS BRAIN Initiative Award 5U19NS104648 (J.W.P.), grant 315230_197296 from the Swiss National Fund (A.P.) and Wellcome Trust grants 209558 (the IBL) and 216324 (the IBL). A.E.U. was supported by the German National Academy of Sciences Leopoldina and the International Brain Research Organization. The funders had no role in study design, data collection and analysis, decision to publish or preparation of the manuscript.

## 6 Author Contributions Statement

Conceptualization, Z.C.A. and J.W.P.; Methodology, Z.C.A. and J.W.P.; Additional Technical and Analysis Support, N.A.R., I.R.S., A.E.U., A.K.C., A.P.; Implementation, Z.C.A.; Data Collection and Curation, A.E.U., A.K.C. and the IBL; Writing – Original Draft, Z.C.A. and J.W.P.; Writing – Review & Editing, Z.C.A., N.A.R., I.R.S., the IBL, A.E.U., A.K.C., A.P. and J.W.P.; Visualization, Z.C.A. and J.W.P.; Supervision, A.P. and J.W.P.; Project Administration, Z.C.A. and J.W.P.; Funding Acquisition, the IBL, A.E.U., A.K.C., A.P. and J.W.P.

## 8 Figures

### 8.1 Main Article Figures

### 8.2 Extended Data Figures

**Figure ED1:**
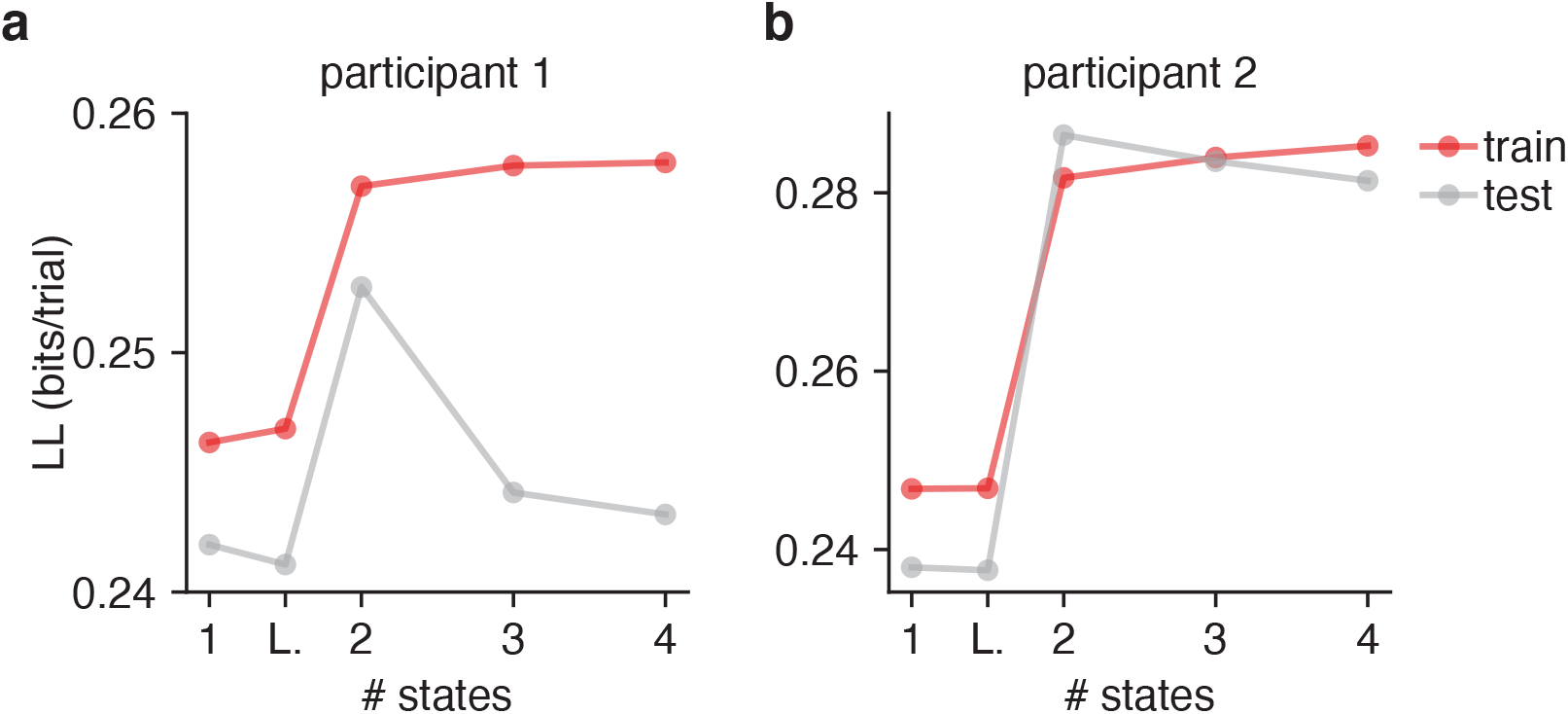
Cross-validation for Model Selection. **(a**,**b)** Here we show the cross-validated train and test loglikelihood in units of bits per trial for two humans performing the task of [27] as a function of the number of states. While the number of parameters of the GLM-HMM increases as the number of states is increased (and a two state GLM-HMM has more parameters than both the classic lapse model and single state GLM), only the loglikelihood of the training dataset (red) is guaranteed to increase as the number of parameters increases. Indeed, as the number of states increases, the GLM-HMM may start to overfit the training dataset causing it to fit the test dataset poorly. This is what we see here when the grey curves begin to decrease in each of the two figures as the number of states increases. Thus, by observing the performance of the model on a test dataset, we can appropriately trade off predictive performance with model complexity.

**Figure ED2:**
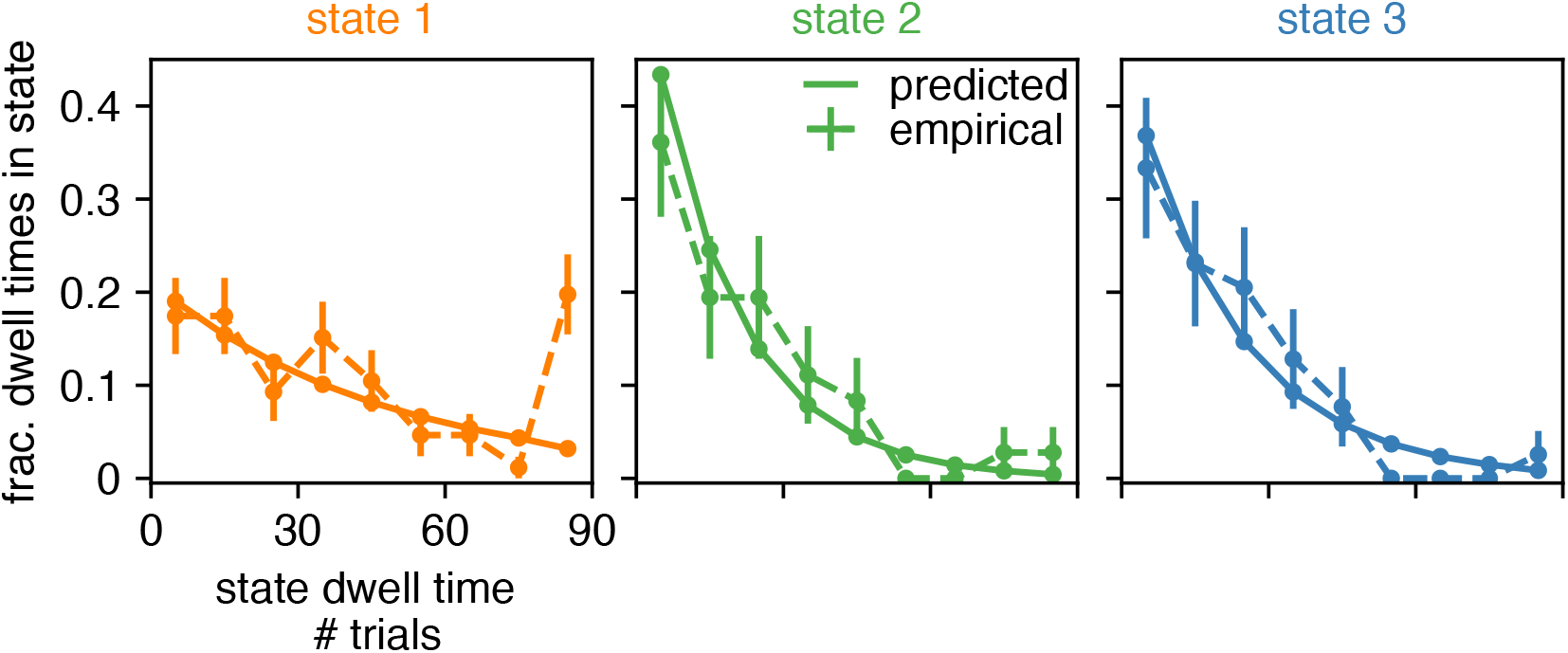
Retrieved state dwell times are approximately geometrically distributed. With the solid line, we show the predicted dwell times (according to the retrieved transition matrix) in each of the three states for the example animal of Fig. 2 and Fig. 3. Predicted dwell times can be obtained from the transition matrix as 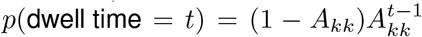 because state dwell times in the Hidden Markov Model are geometrically distributed. We then use the posterior state probabilities to assign states to trials in order to calculate the dwell times that are actually observed in the real data (shown with the dashed line); we also show the 68% confidence intervals associated with these empirical probabilities (n is between 36 and 86, depending on state). We find that the empirical dwell times for the biased leftward and rightward states seem to be geometrically distributed. For the engaged state, because there are some entire sessions (each session is 90 trials) during which the animal is engaged, we see that the empirical dwell times associated with this state are not as well described by a geometric distribution. A future direction may be to allow non-geometrically distributed state dwell times by replacing the Hidden Markov Model with e.g. the Hidden semi-Markov Model [0].

**Figure ED3:**
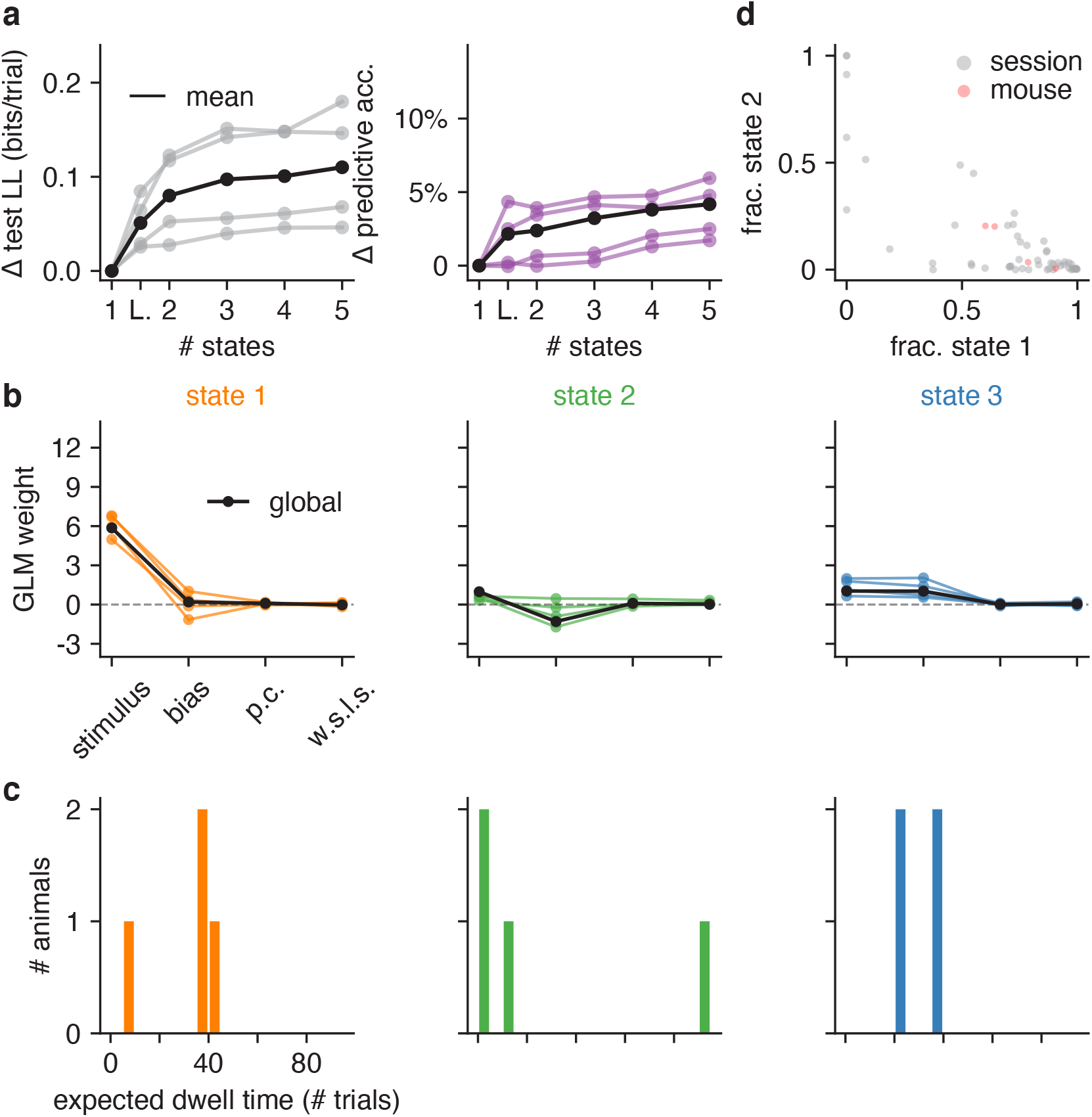
GLM-HMM application to 4 mice not exposed to bias blocks in IBL task. We confirm that mice that have never been exposed to bias blocks in the IBL task continue to show state-dependent decision-making. This is a sister figure to Fig. 4, and each panel can be interpreted in the same way as in Fig. 4.

**Figure ED4:**
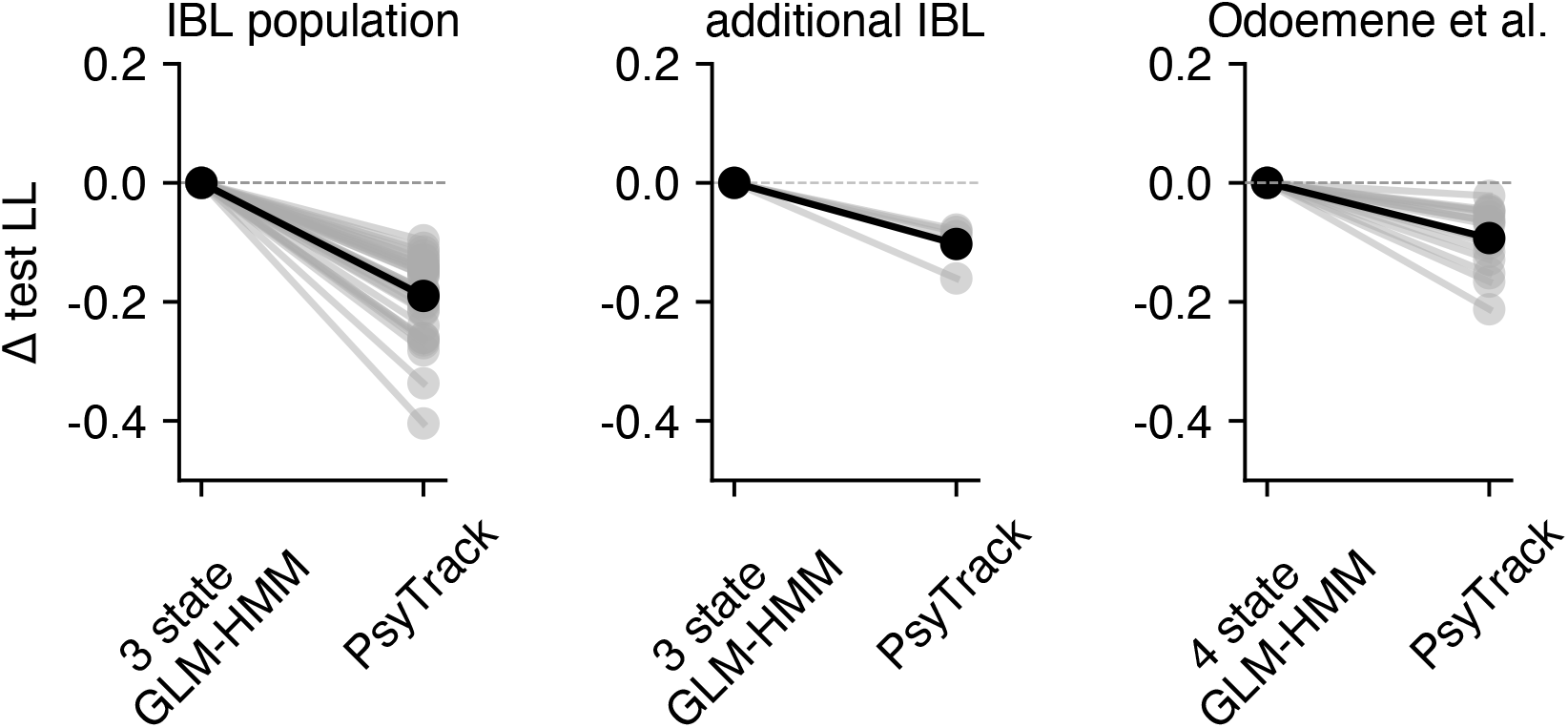
Additional comparisons with PsyTrack model. **(a)** Copy of left panel from Fig. 4f showing the difference in test set loglikelihood for 37 IBL animals for the 3 state GLM-HMM compared to the PsyTrack model of [35, 36]. Black indicates the mean across animals. The 3 state GLM-HMM better explained the choice data of all 37 animals compared to the PsyTrack model with continuously evolving states. **(b)** Analogous figure for the 4 additional IBL animals studied in Fig. ED3. All 4 animals’ data were better explained by the GLM-HMM compared to PsyTrack. **(c)** Same as in panel a for Odoemene et al. animals shown in Fig. 5, although comparison now utilizes 4 state GLM-HMM fits. All 15 animals’ data were better explained by the GLM-HMM compared to PsyTrack.

**Figure ED5:**
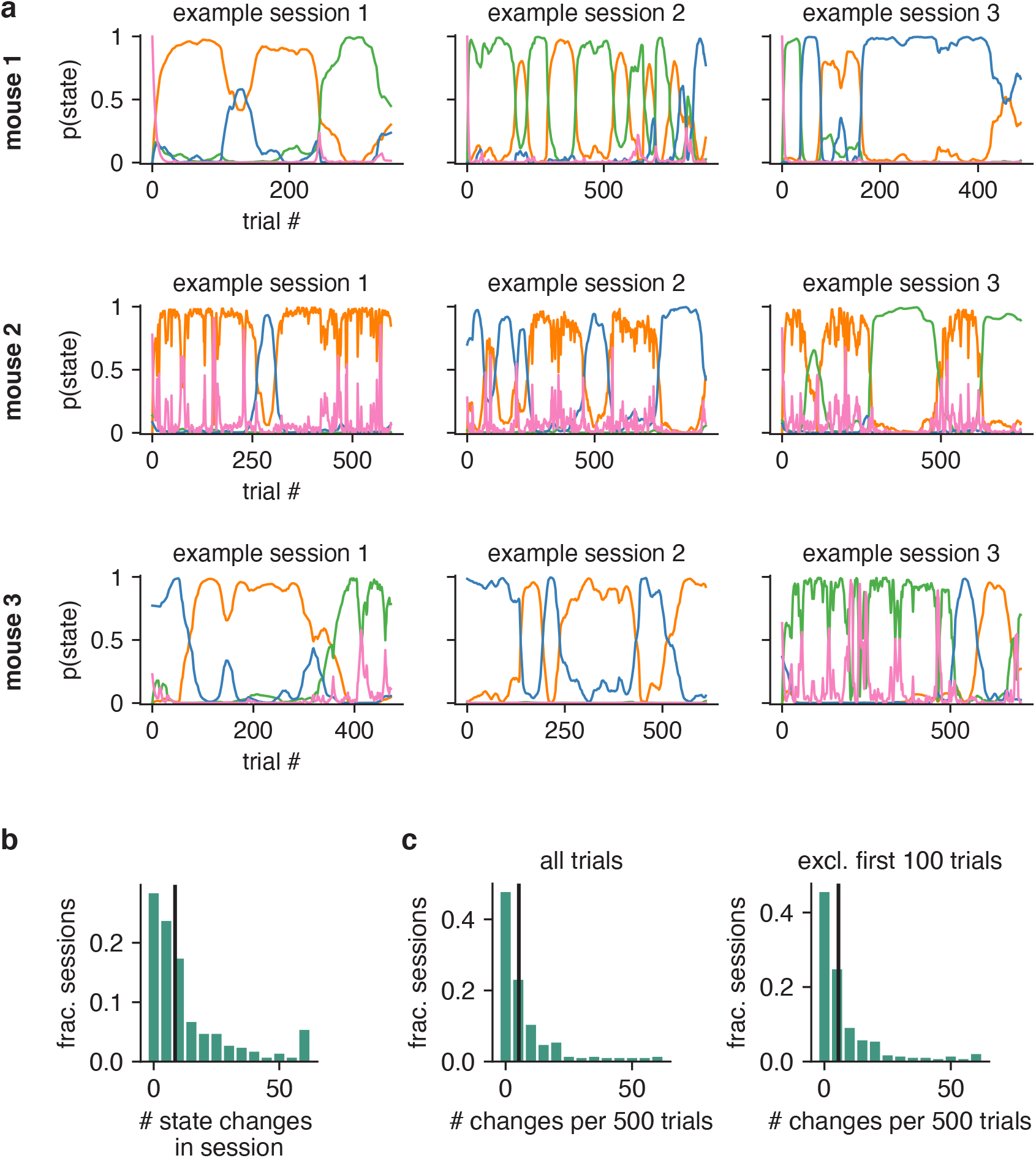
State switching in Odoemene et al. task. **(a)** Posterior state probabilities for three example sessions for three example mice (different mice are shown in each row). **(b)** Histogram giving number of state changes (identified with posterior state probabilities) per session for all sessions across all animals. For visibility, state changes are censored above 60. **(c)** Different sessions have different numbers of trials, so we normalize the histogram of b to give the number of state changes per 500 trials for each session (the median session length is 683 trials). Again, for visibility, state changes are censored above 60. Left: we use all data from all trials to generate the normalized histogram. Right: we plot the normalized histogram when we exclude the first 100 trials of a session. As can be observed, the left and right normalized histograms are very similar (p-value = 0.96 using KS-test). While the GLM-HMM is able to capture “warm-up” effects (as described in the main text), this test reveals that the GLM-HMM is able to capture more than this, and state switching occurs much later in the session too (as also indicated by the posterior state probabilities shown in a).

**Figure ED6:**
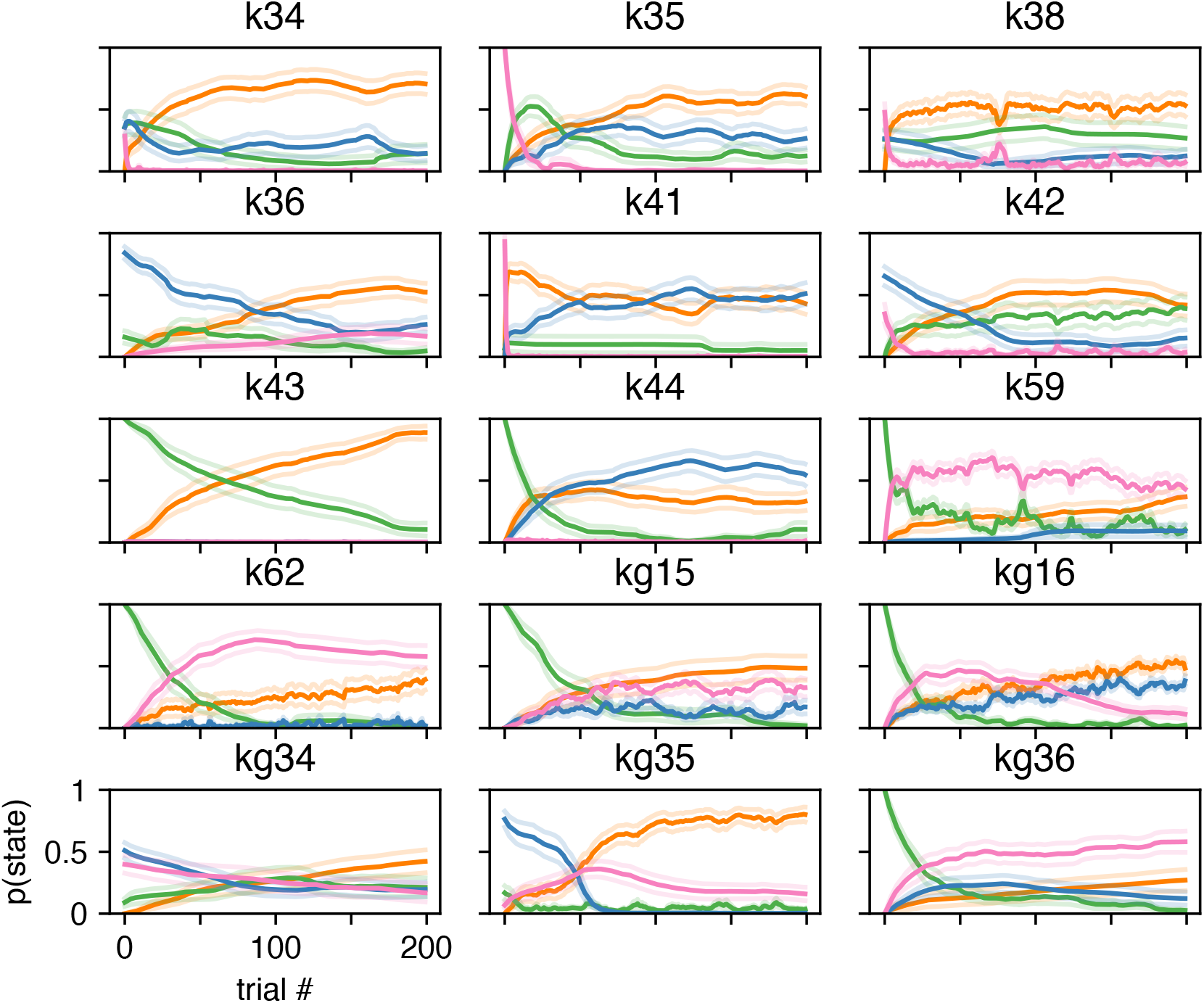
GLM-HMM captures ‘warm-up’ effect for Odoemene et al. animals. Average (across 20 sessions) posterior state probabilities for the first 200 trials of a session for each animal in the Odoemene et al. dataset. Orange corresponds to the engaged state, green to the biased left, blue to the biased right and pink to the win-stay state from Fig. 5. Error bars represent standard errors.

**Figure ED7:**
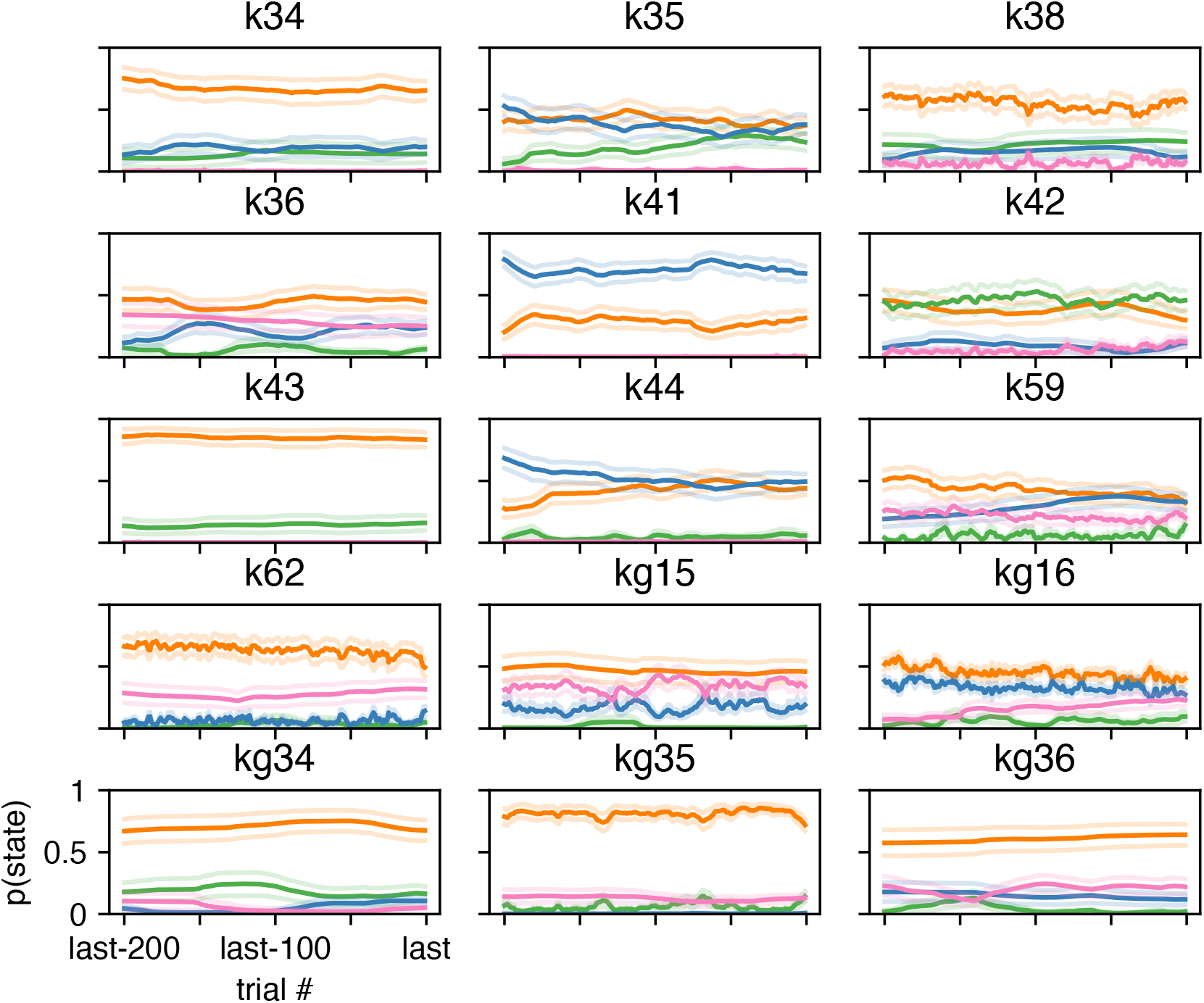
GLM-HMM posterior state probabilities at end of session. Average (across 20 sessions) posterior state probabilities for the last 200 trials of a session for each animal in the Odoemene et al. dataset. Orange corresponds to the engaged state, green to the biased left, blue to the biased right and pink to the win-stay state from Fig. 5. Error bars represent standard errors.

**Figure ED8:**
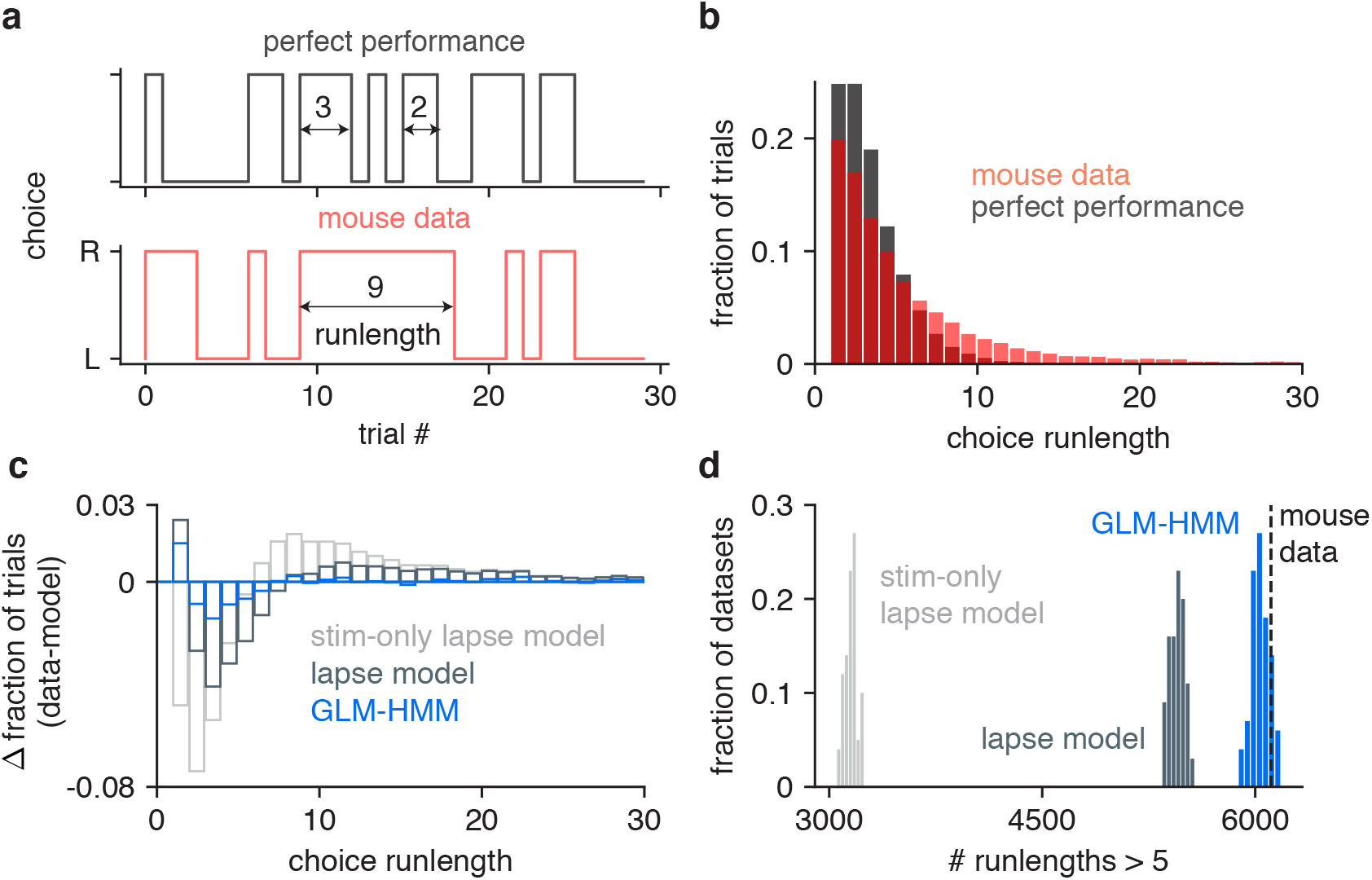
Simulated data from GLM-HMM captures statistics of real choice data. **(a)** Definition of choice run-length. Shown are the choices that an IBL mouse made over the course of 30 trials (red, bottom), as well as the choices it should have made during that same time course if the mouse performed the task perfectly (grey, top). Choice run-length is defined as the number of trials during which a mouse repeated the same decision (example choice run-lengths of 2, 3 and 9 trials are highlighted). **(b)** Red: fraction of trials in choice run-lengths of between 1 and 30 trials when calculated from all trials for all mice. Grey: distribution of choice run-lengths that would have been obtained if IBL mice performed the task perfectly. **(c)** Difference in choice run-length distribution for simulated data (from three different models) compared to the red distribution shown in (b). Models used to simulate data were a lapse model with only stimulus intensity and bias regressors, a lapse model that also included history regressors (previous choice and win-stay-lose-switch), and a 3 state GLM-HMM (also with history regressors). We simulated 100 example choice sequences from each model and calculated the mean histogram of choice run-lengths across the 100 simulations. This was then subtracted from the red histogram of (b). **(d)** Number of choice run-lengths with more than 5 trials for each model simulation used in (c). In the 181,530 trials of real choice data, there were 6111 run-lengths lasting more than 5 trials (as shown with the dashed line). When we simulated choice data according to each of the models shown in (c), we found that only the GLM-HMM could generate simulations with as many run-lengths lasting more than 5 trials as in the real data (15/100 simulations had 6111 or more run-lengths lasting more than 5 trials for the GLM-HMM compared to 0/100 for both of the lapse models).

**Figure ED9:**
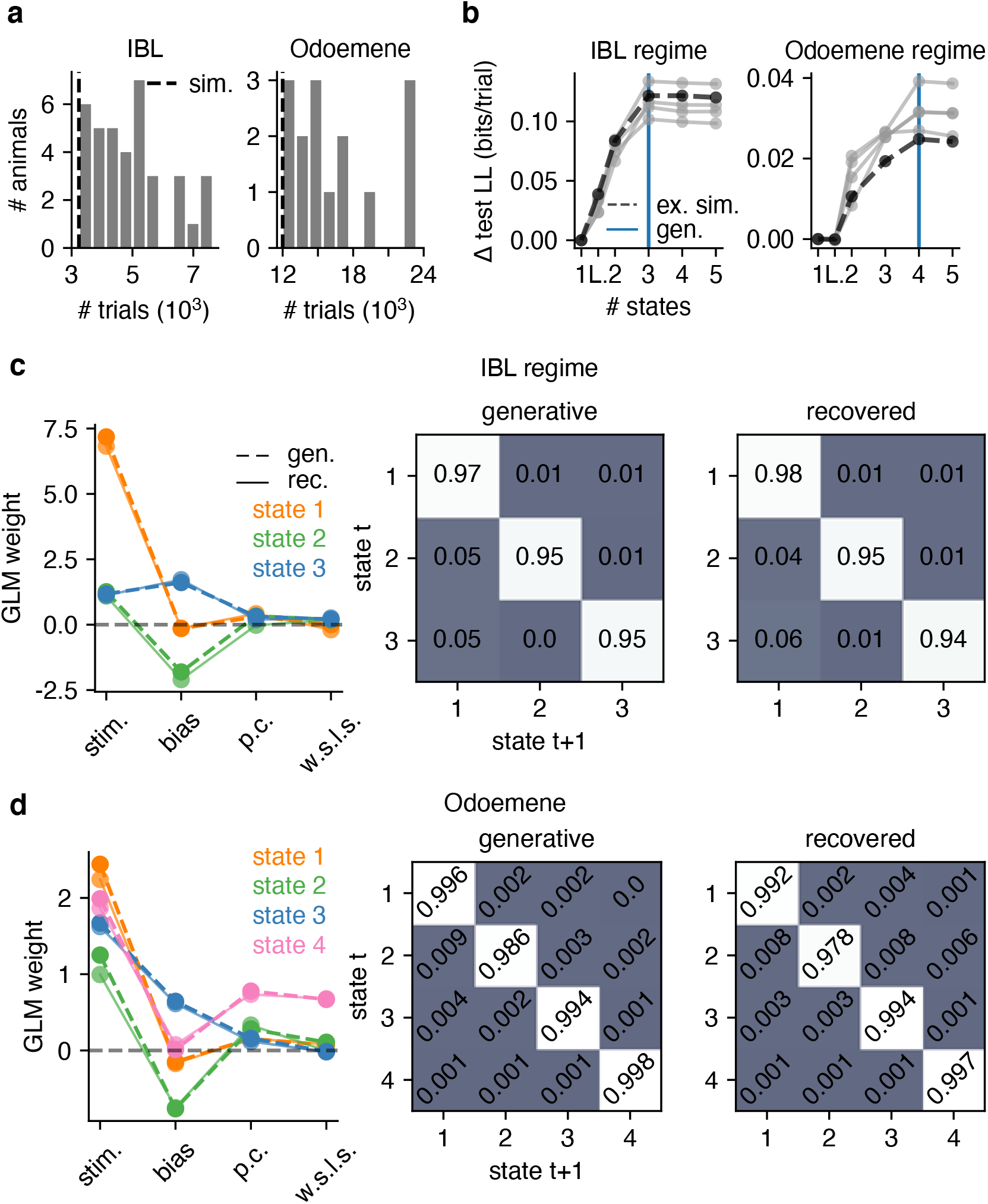
GLM-HMM Recovery Analysis 1. For dataset sizes comparable to those of real animals, we can recover the IBL and Odoemene global parameters in simulated data. **(a)** Dataset sizes for each of the 37 IBL animals studied (left) and 15 mice from Odoemene et al. (right). The dashed vertical line indicates the number of trials that we used in simulation data in panels b, c and d (3240 for the IBL parameter regime and 12000 for the Odoemene regime simulation). **(b)** Test set loglikelihood for each of 5 simulations is maximized at 3 states (blue vertical line) after we simulate according to the (IBL regime) parameters shown in panel c. Similarly, in the right panel, test set loglikelihood is maximized at 4 states when we simulate choice data with the (Odoemene regime) parameters shown in panel d. The thick black line marked as ‘ex. sim.’ (example simulation) indicates the simulation whose generative and recovered parameters we show in panels c and d. **(c)** Left: the generative and recovered GLM weights (for the simulation marked as ‘ex. sim.’ in panel b) when we simulate choice data in the IBL parameter regime. Middle and right: the generative and recovered transition matrices. **(d)** The generative and recovered parameters in the Odoemene et al. parameter regime.

**Figure ED10:**
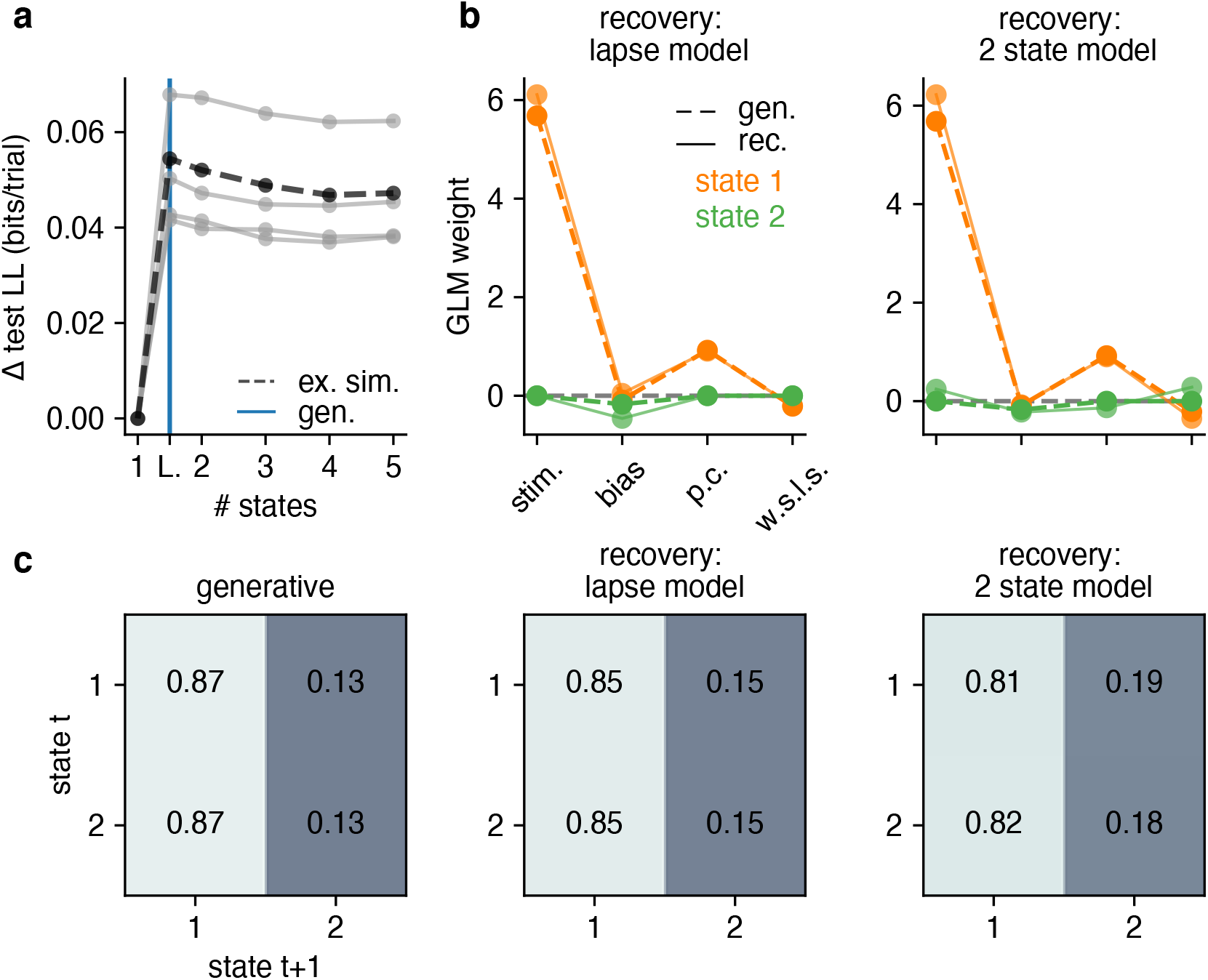
GLM-HMM Recovery Analysis 2: We can recover lapse behavior. **(a)** We simulate 5 datasets, each with 3240 trials, according to the best fitting lapse model for IBL animals. We then fit these simulated datasets with GLM-HMMs, as well as a lapse model (a constrained 2 state GLM-HMM). The test set loglikelihood is highest for the lapse model in all simulations, indicating that lapse behavior can be distinguished from the long-enduring multi-state behavior that best described the real data. The thick black line marked as ‘ex. sim.’ (example simulation) indicates the simulation whose generative and recovered parameters we show in panels b and c. **(b)** Left: the generative and recovered weights when recovery is with a lapse model. Right: the generative weights are the same as in the left panel, but we now recover with an unconstrained 2 state GLM-HMM (thus the stimulus, previous choice and w.s.l.s. weights for the second state can be non-zero) **(c)** The generative (left) transition matrix and the recovered transition matrices when we recover with a lapse model (middle) and an unconstrained 2 state GLM-HMM (right). While the lapse model and 2 state GLM-HMM results don’t perfectly agree, if mice were truly lapsing, the transition matrix would not have the large entries on the diagonals that we observe in the real data.

## Supplementary Information

**Figure S1:**
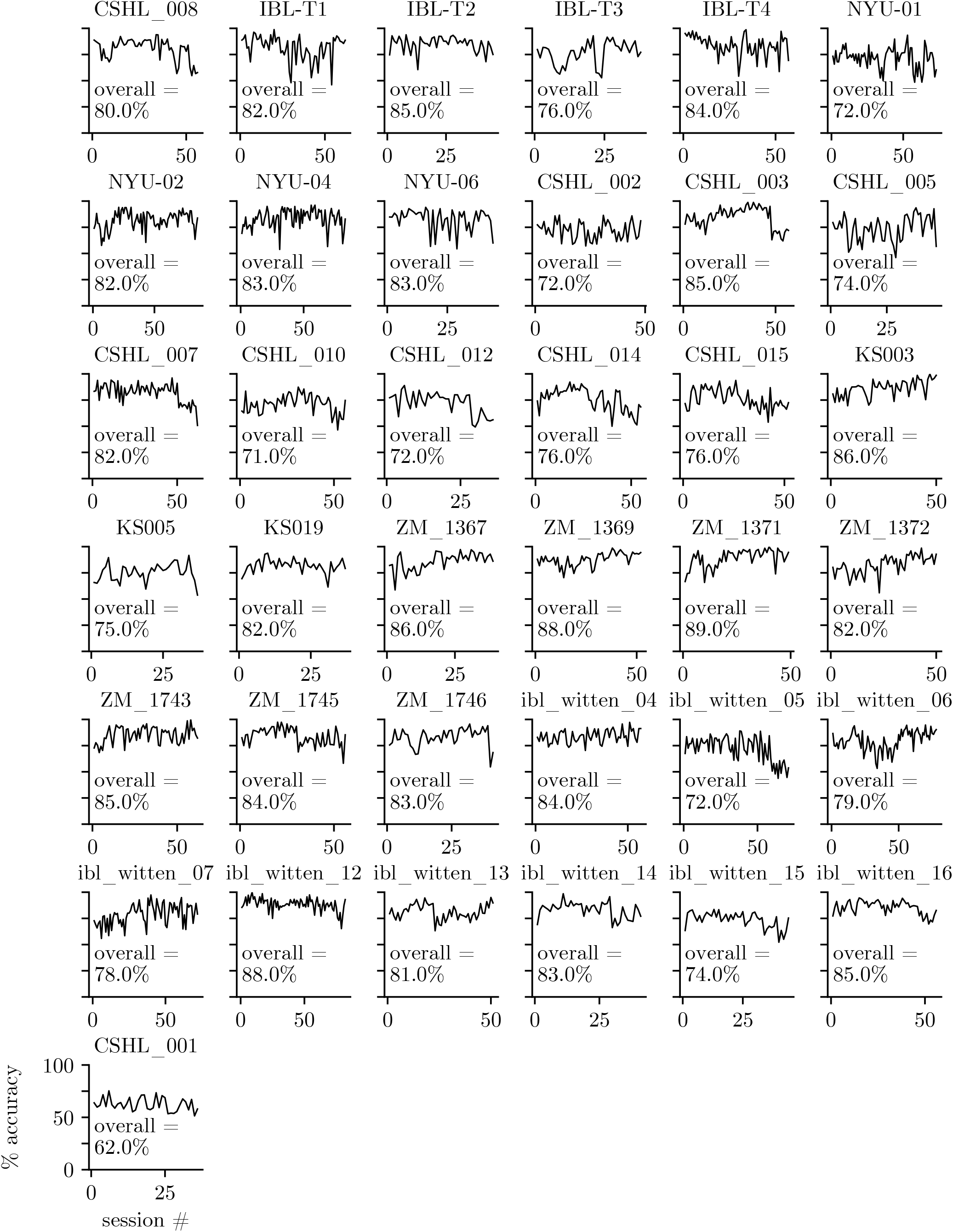
Raw behavioral data for IBL animals: Accuracy across sessions. We plot the accuracy of IBL animals [19] across sessions as evidence that mice have learned the task, and that their choice behavior has reached stationarity. We also report the overall accuracy of each animal when aggregated across sessions. The example animal studied in Fig. 2 and Fig. 3 is ‘CSHL_008’.

**Figure S2:**
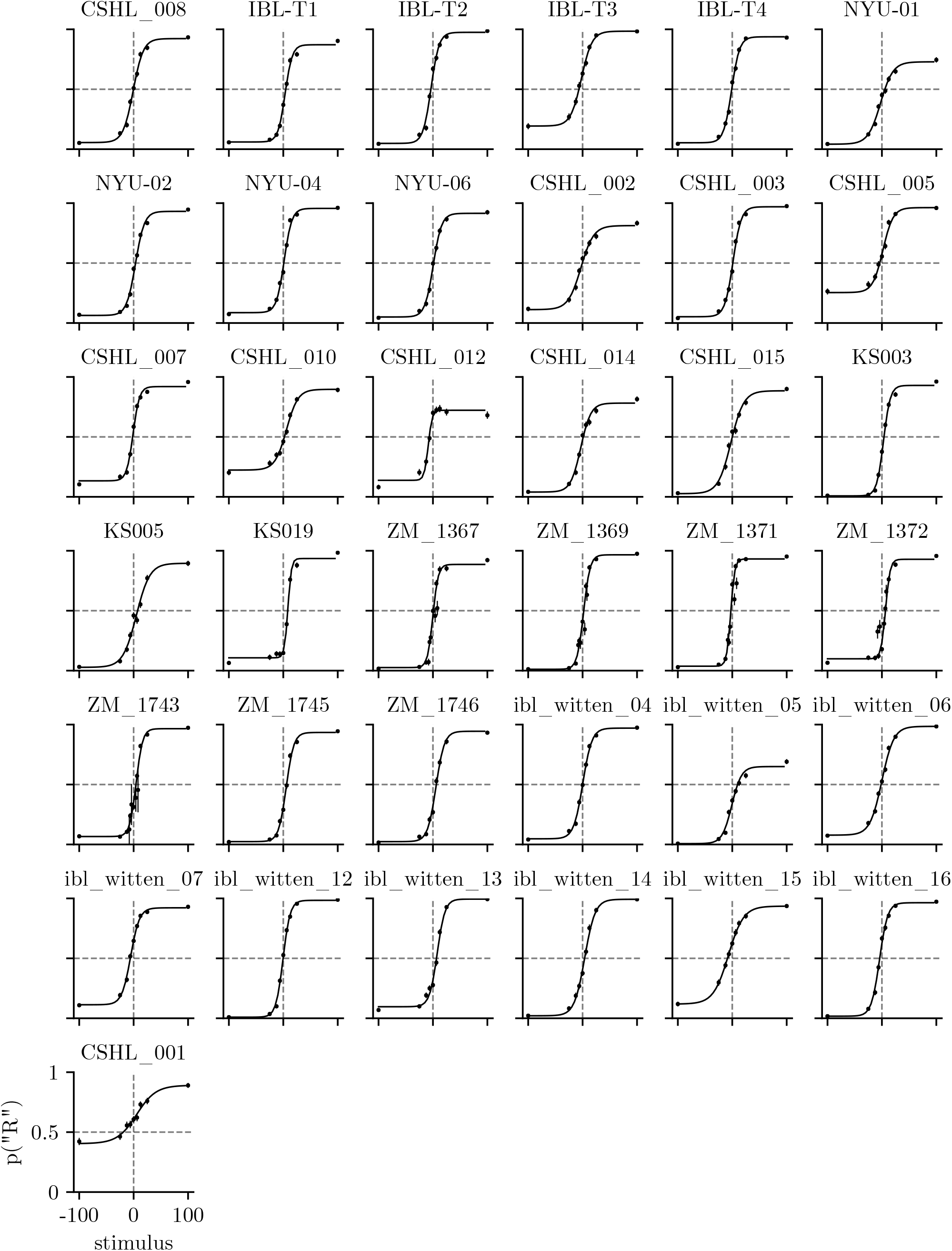
Raw behavioral data for IBL animals: Psychometric curves. We plot the psychometric curves for the 37 IBL animals [19] whose choice data we study. The example animal that we study in Fig. 2 and Fig. 3 is ‘CSHL_008’. Animals are ordered in the same way that they are in Fig. S1 when considering row-major order, so plots can be compared across the two figures. In each plot, we also show each animal’s empirical choice probabilities; error bars are 68% confidence intervals (minimum n for animal-stimulus pair was 6, while maximum was 1547. Median n was 493).

**Figure S3:**
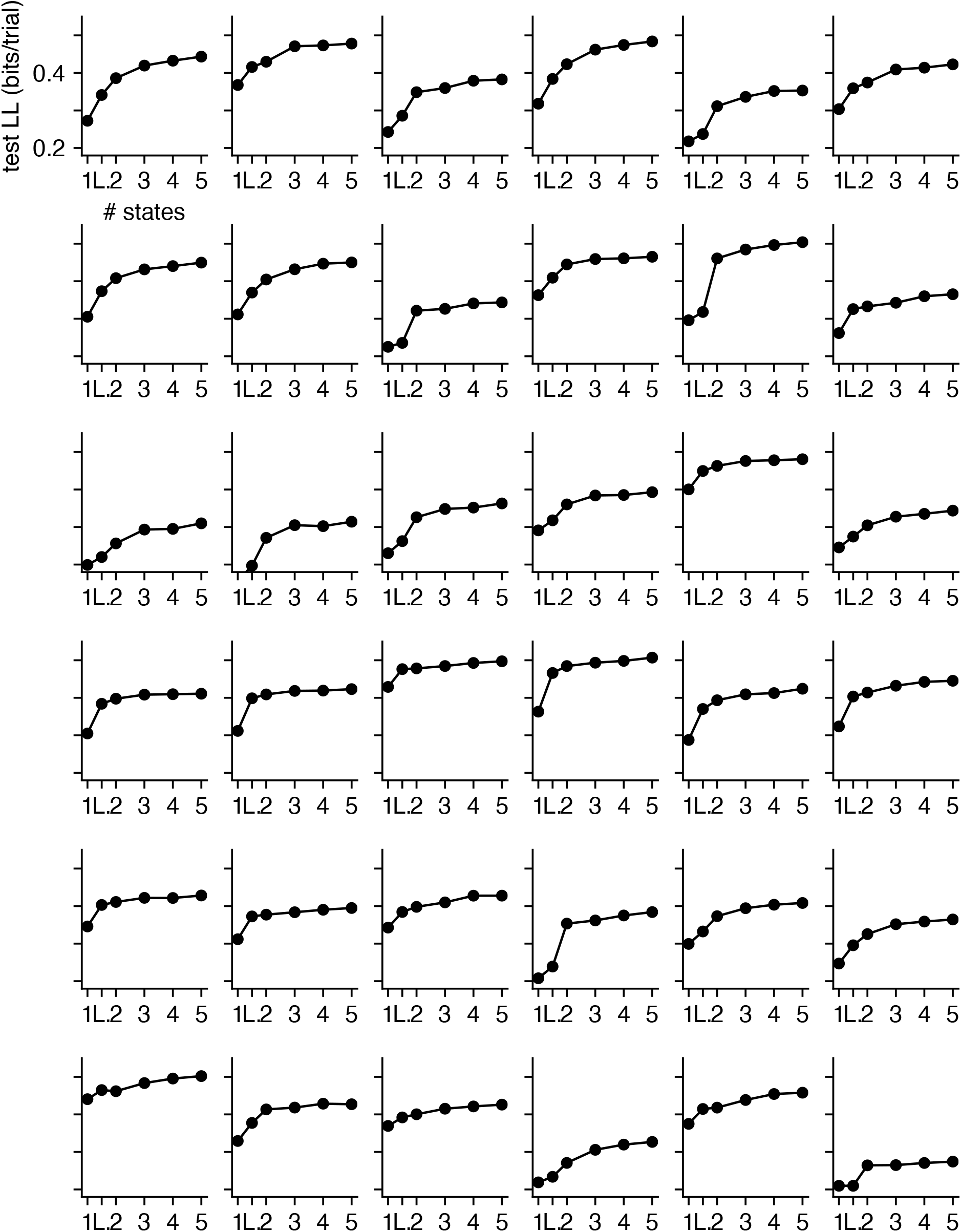
Model comparison for all IBL animals. Rather than plotting curves on top of one another as in Fig. 4a, we plot each individual animal’s test set loglikelihood in a grid. We do not show model comparison results for the example animal studied in Fig. 2 and Fig. 3 since we show the curve for that particular animal there, but animals are otherwise ordered in the same way (according to row-major order) that they are in Fig. S1 and Fig. S2, so plots can be compared across these figures.

**Figure S4:**
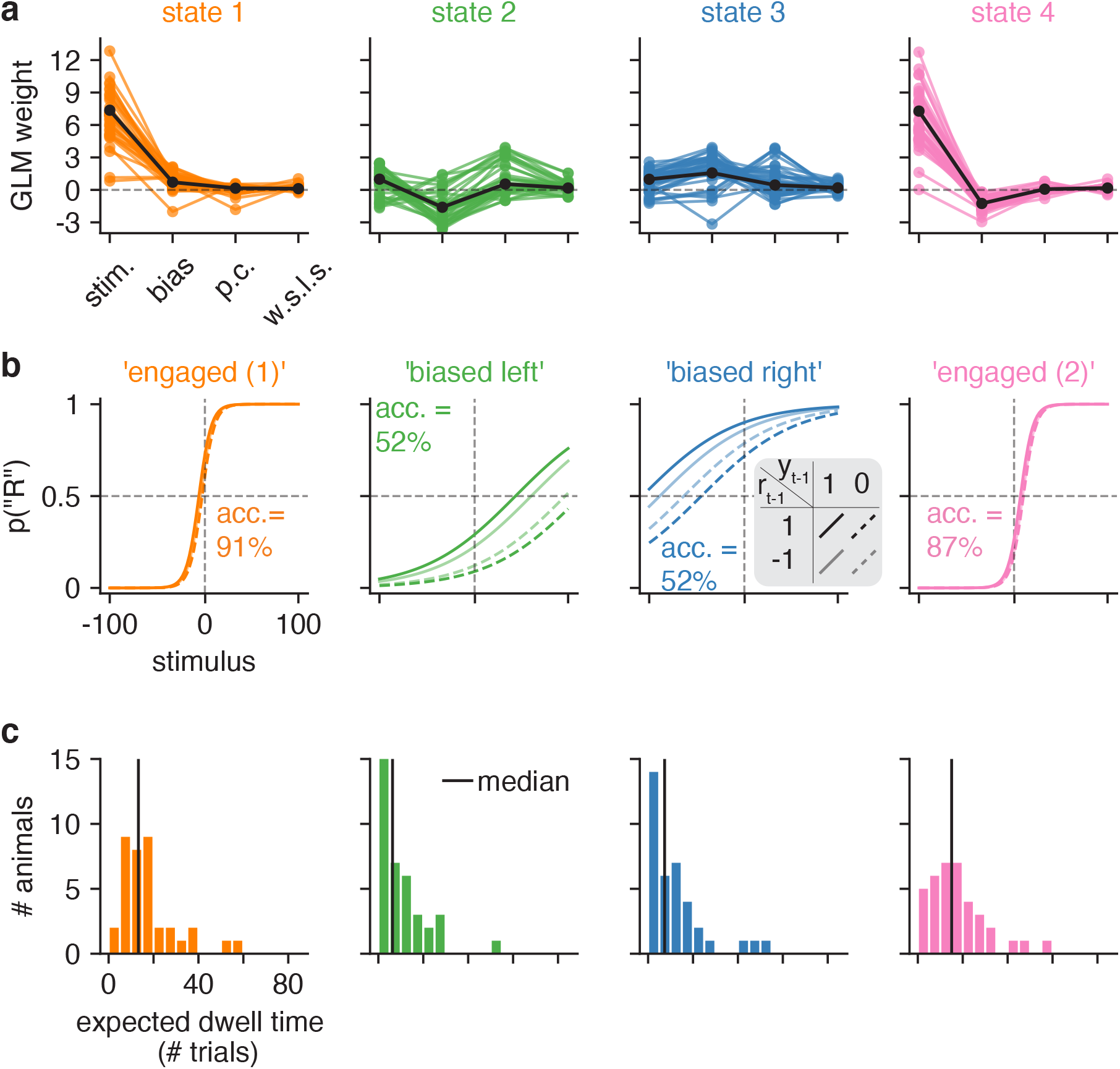
GLM-HMM 4 state fits to IBL data. **(a)** Retrieved GLM weights for each of the 37 IBL animals for each state of the 4 state GLM-HMM, as well as the global weights (black) for the fit when all data from all animals are concatenated. **(b)** An alternative method of plotting the weights in panel a: we plot, as a function of the stimulus intensity, as well as the animal’s reward and choice on the previous trial, the probability that the animal goes rightward at the current trial. **(c)** The expected dwell time for each animal in each of the 4 states, as obtained from the best-fitting transition matrices for each animal.

**Figure S5:**
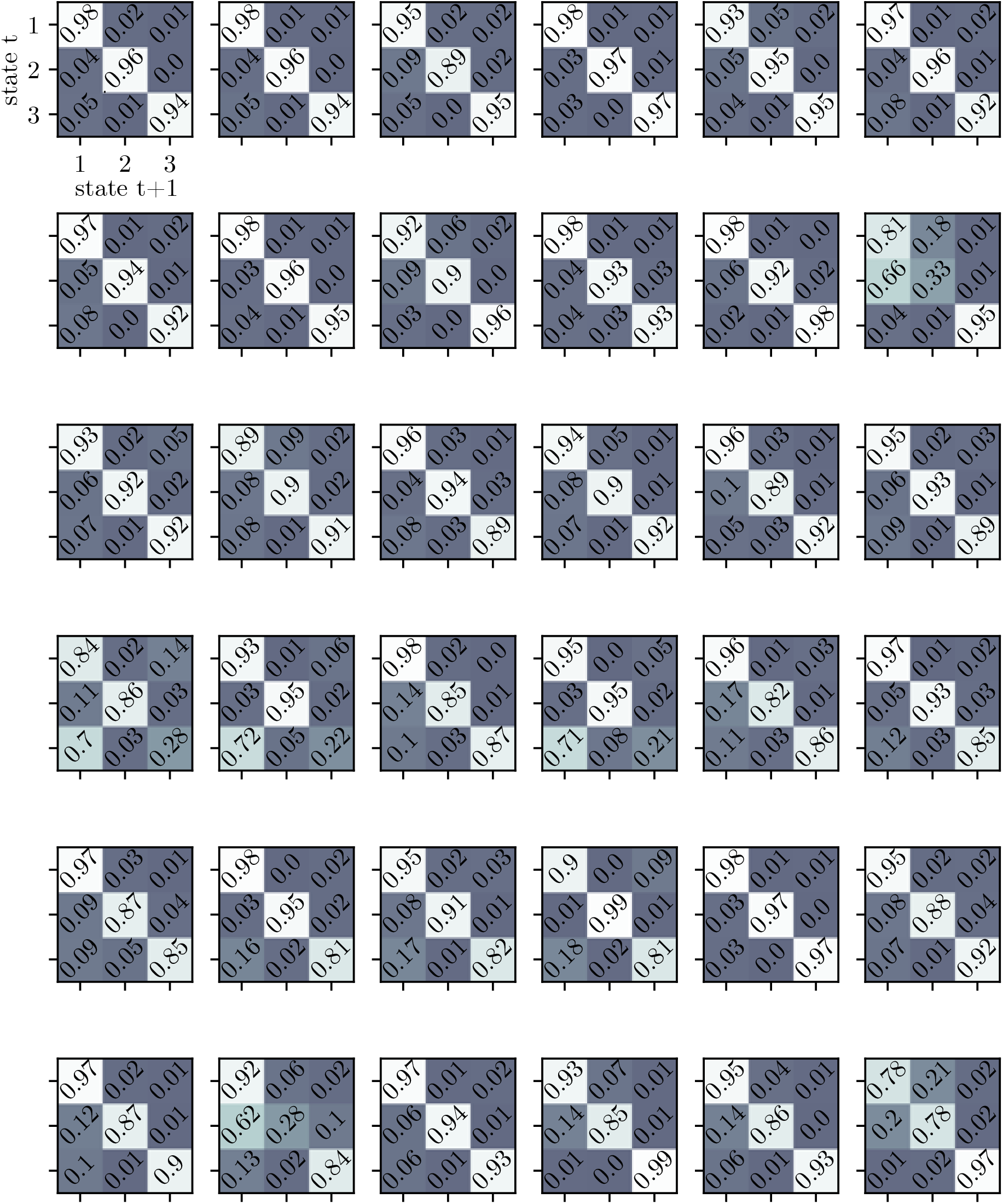
Retrieved transition matrices for all IBL animals. We do not plot the transition matrix for the example animal studied in Fig. 2 as the retrieved transition matrix is shown there. Animals are ordered in the same way as in other supplemental figures so plots can be compared across figures.

**Figure S6:**
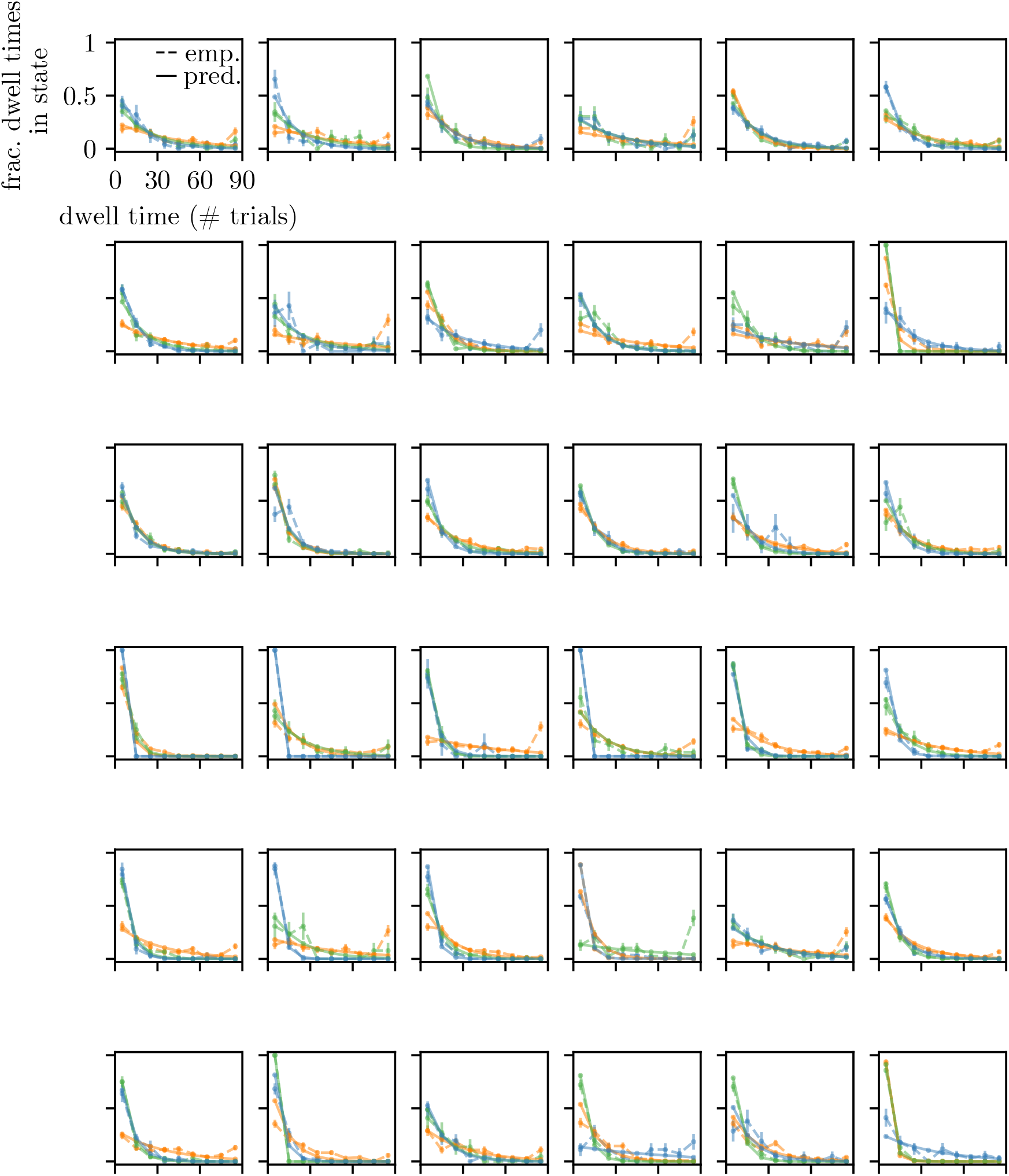
State dwell times are approximately geometrically distributed for all IBL animals. We plot the predicted (“pred.”; shown with the solid line) and empirical (“emp.”; shown with the dashed lines) state dwell time probabilities for each IBL animal and each state (excluding the example animal since the state dwell time probabilities for the example animal are shown in Fig. ED2)). Colors map to states in the usual way (orange is state 1, green is state 2, blue is state 3). The empirical state dwell times are obtained by using the posterior state probabilities to assign state labels to trials. 68% confidence intervals are shown for empirical values (median n for an animal-state pair was 75; minimum n was 9; maximum n was 419). Predicted state dwell time probabilities are obtained from the transition matrix according to 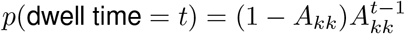.

**Figure S7:**
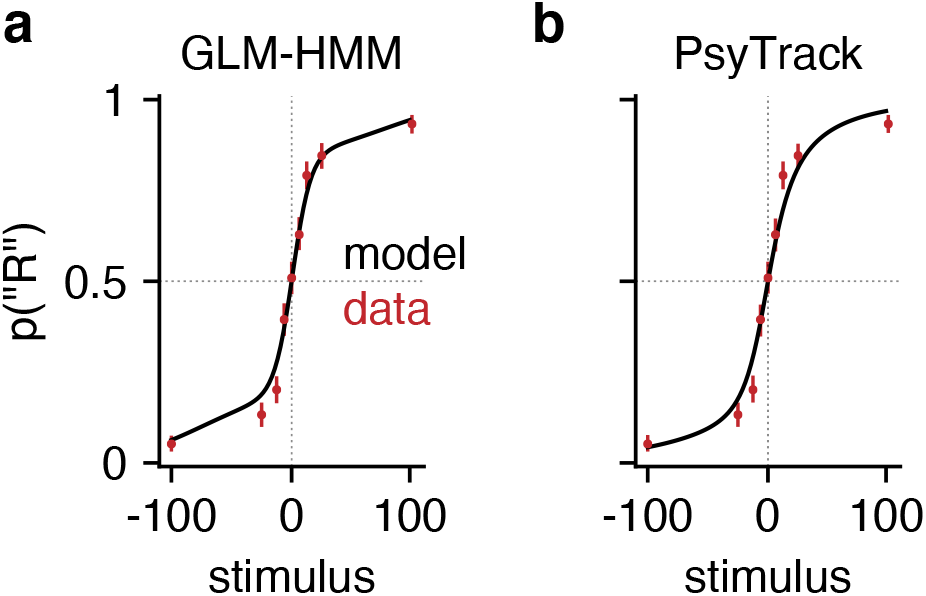
GLM-HMM and PsyTrack model fits to choice data from example mouse. **(a)** Copy of Fig. 2h illustrating the psychometric curve generated by the GLM-HMM for the example mouse studied in Fig. 2 and Fig. 3. Specifically, using the fit GLM-HMM parameters for this animal and the true sequence of stimuli presented to the mouse, we generated a time series with the same number of trials as those that the example mouse had in its dataset. At each trial, regardless of the true stimulus presented, we calculated *p*_*t*_(“*R*”) for each of the 9 possible stimuli by averaging the per-state psychometric curves of Fig. 2g and weighting by the appropriate row in the transition matrix (depending on the sampled latent state at the previous trial). Finally, we averaged the per-trial psychometric curves across all trials to obtain the curve that is shown in black, while the empirical choice data of the mouse are shown in red, as are 95% confidence intervals (n between 530 and 601 depending on stimulus). **(b)** Ability of PsyTrack model to fit the empirical choice data (red dots with 95% confidence intervals (n between 530 and 601 depending on stimulus)) for the same example mouse. We again obtained a per-trial psychometric curve using the per-trial weights returned by the PsyTrack model of [35, 36] and used these to evaluate *p*_*t*_(“*R*”) for each of the 9 possible stimuli. Again, the black line represents the average curve across all trials. Note: despite the absence of explicit lapse parameters in this model, it is able to capture the non-zero error rate of the mouse on “easy” trials.

**Figure S8:**
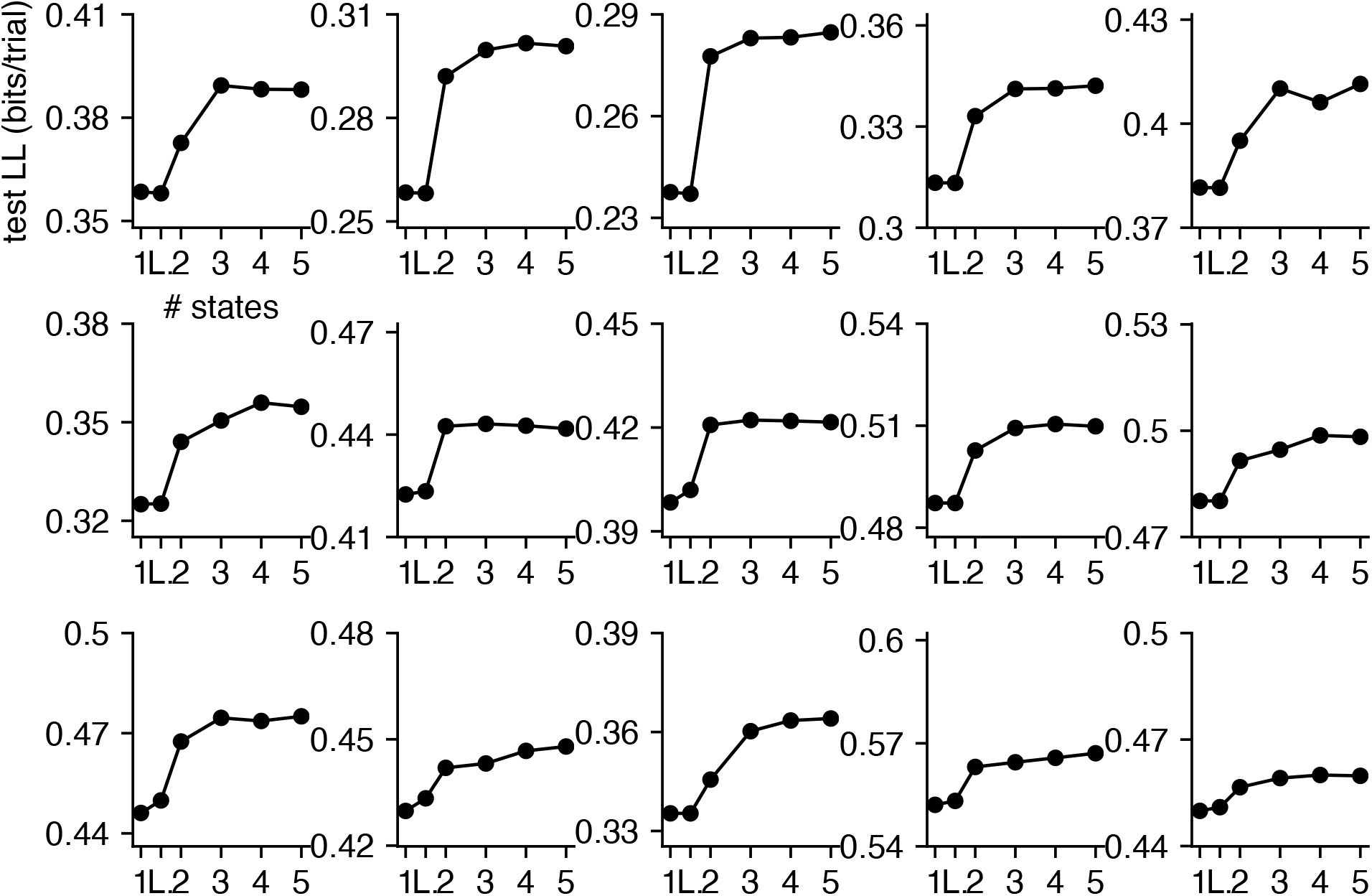
Model Comparison for all Odoemene et al. animals. Rather than plotting curves on top of one another as in Fig. 5b, we plot each individual animal’s test set loglikelihood in a grid. Animals are ordered in the same way that they are in Fig. S13 and Fig. S14, so plots can be compared across figures.

**Figure S9:**
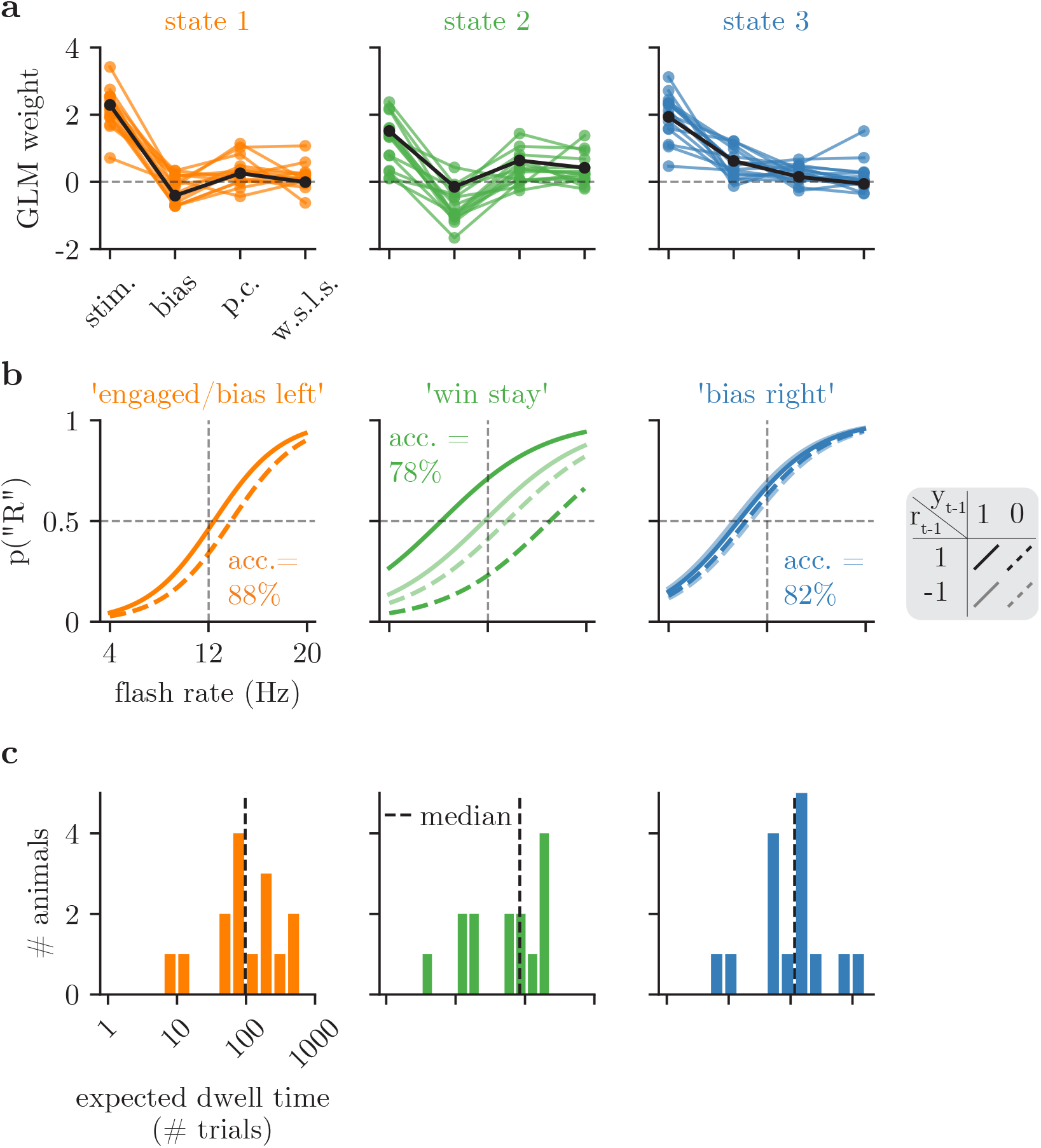
GLM-HMM 3 state fits to Odoemene et al. data. When 3 state GLM-HMMs are fit to the Odoemene et al. data, the engaged and bias left states are merged to form a single (mostly engaged) state (accuracy remains high at 88%), while the win-stay and bias right states are largely unchanged. This is a sister figure to panels d, e, and f of Fig. 5, where the panels can be interpreted in the same way that they are there.

**Figure S10:**
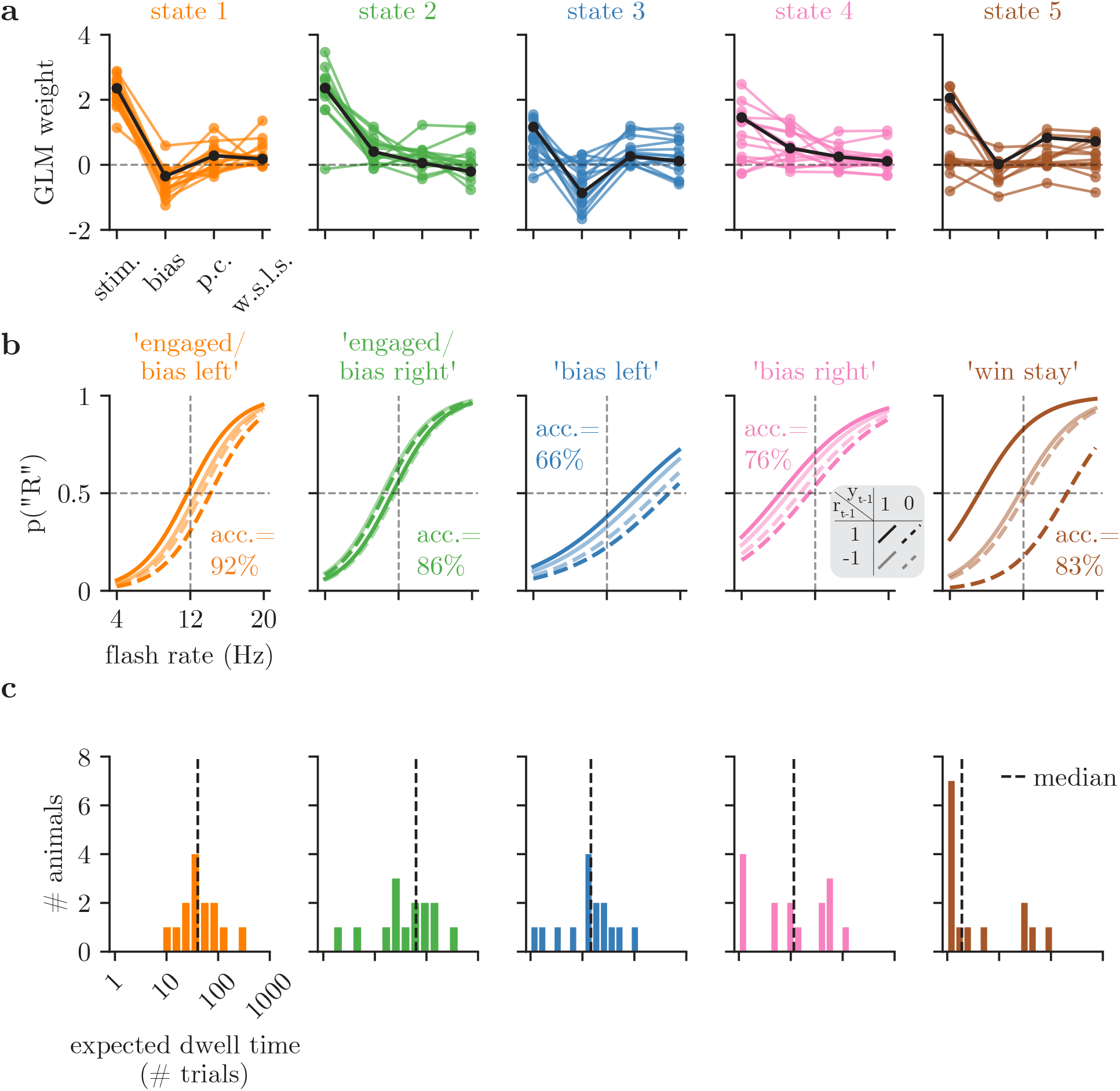
GLM-HMM 5 state fits to Odoemene et al. data. When 5 state GLM-HMMs are fit to the Odoemene et al. data, the engaged state is split into an engaged/bias left and engaged/bias right state, while the win-stay, bias right and bias left states are largely unchanged. This is a sister figure to panels d, e, and f of Fig. 5, where the panels can be interpreted in the same way that they are there.

**Figure S11:**
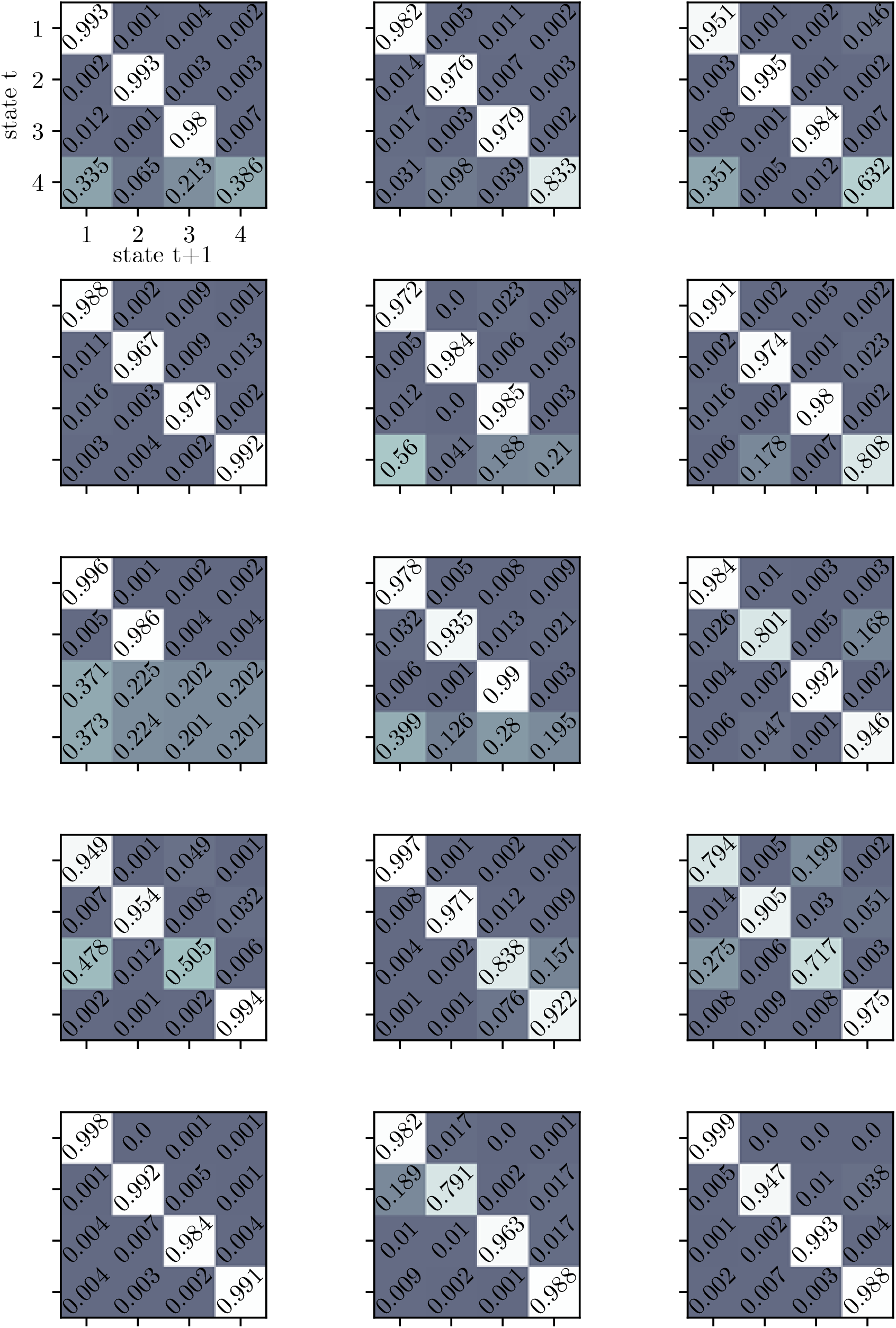
Retrieved transition matrices for Odoemene et al. animals. Each individual transition matrix is the best fitting transition matrix for 1 of the 15 Odoemene et al. mice that we study. Row-major order has animals ordered in the same way that they are in Fig. S11.

**Figure S12:**
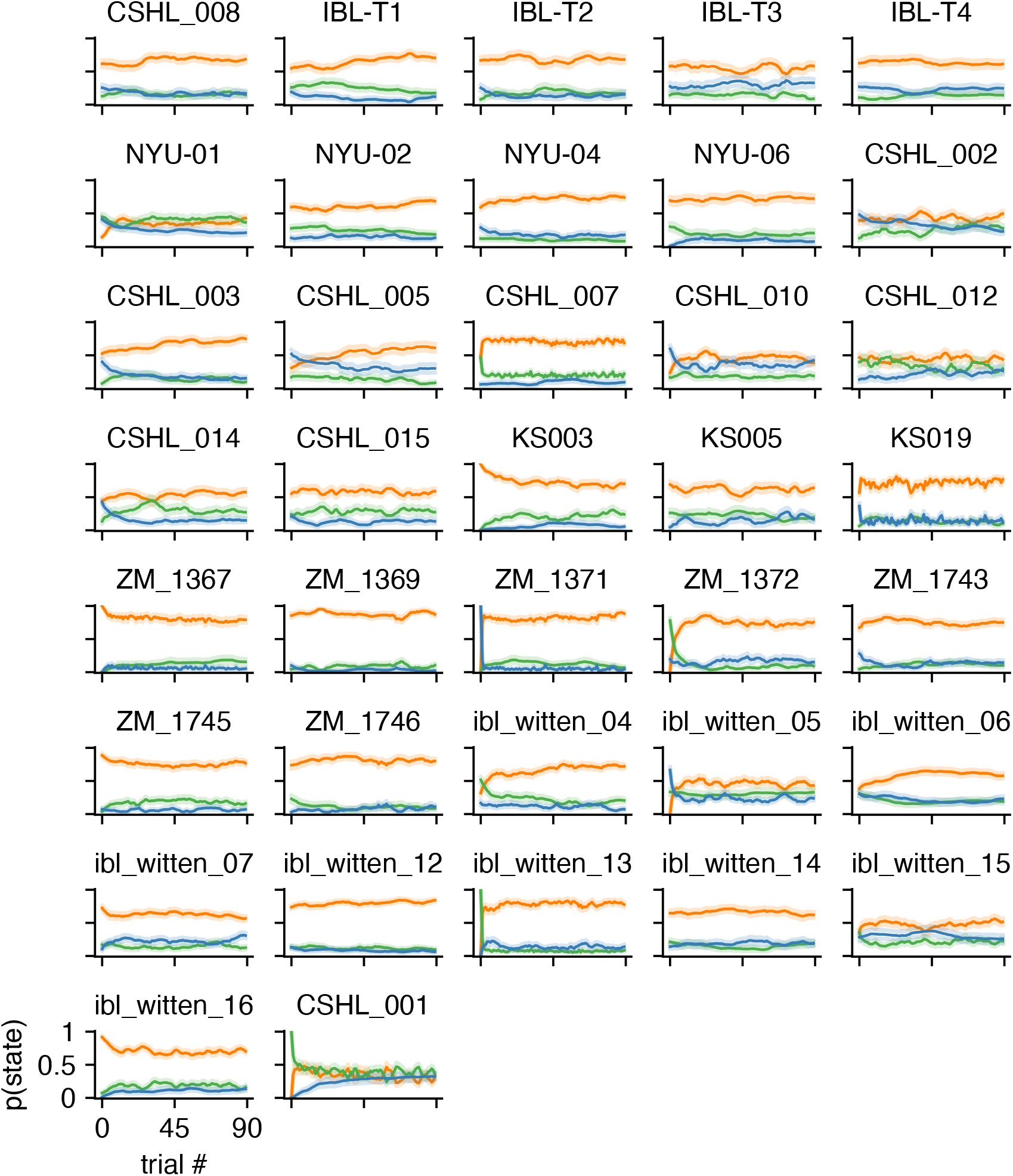
Average posterior state probabilities for all IBL animals. Average posterior state probabilities across all sessions for each individual animal along with 68% confidence intervals (minimum n across animal-state pairs: 36; maximum was n=87). Each session lasts 90 trials. Animals are ordered in the same way as in other supplemental figures so plots can be compared across figures.

**Figure S13:**
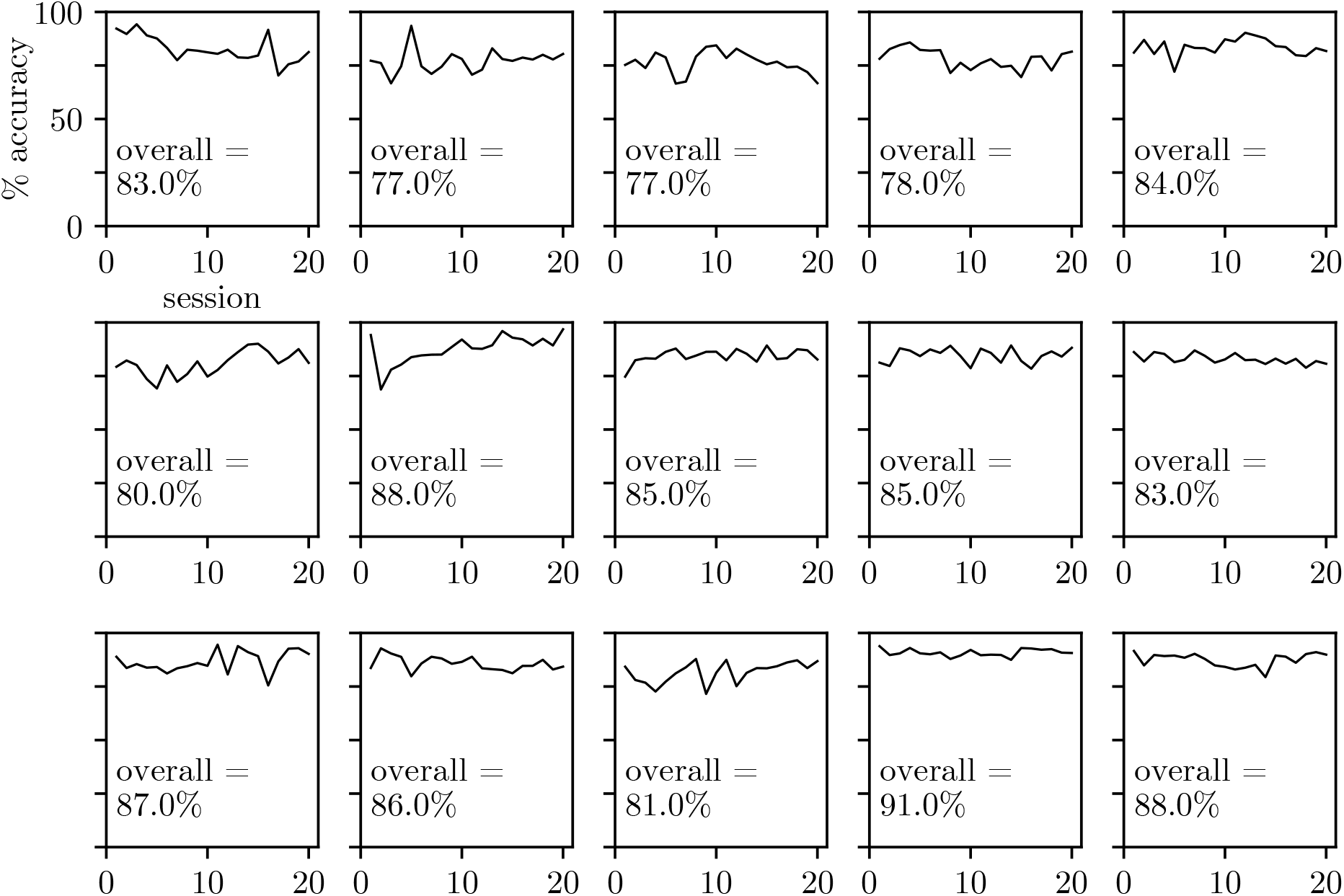
Accuracy across sessions for Odoemene et al. [20] animals. We plot the accuracy across sessions for Odoemene et al. animals as evidence that the animals’ choice behavior has reached stationarity.

**Figure S14:**
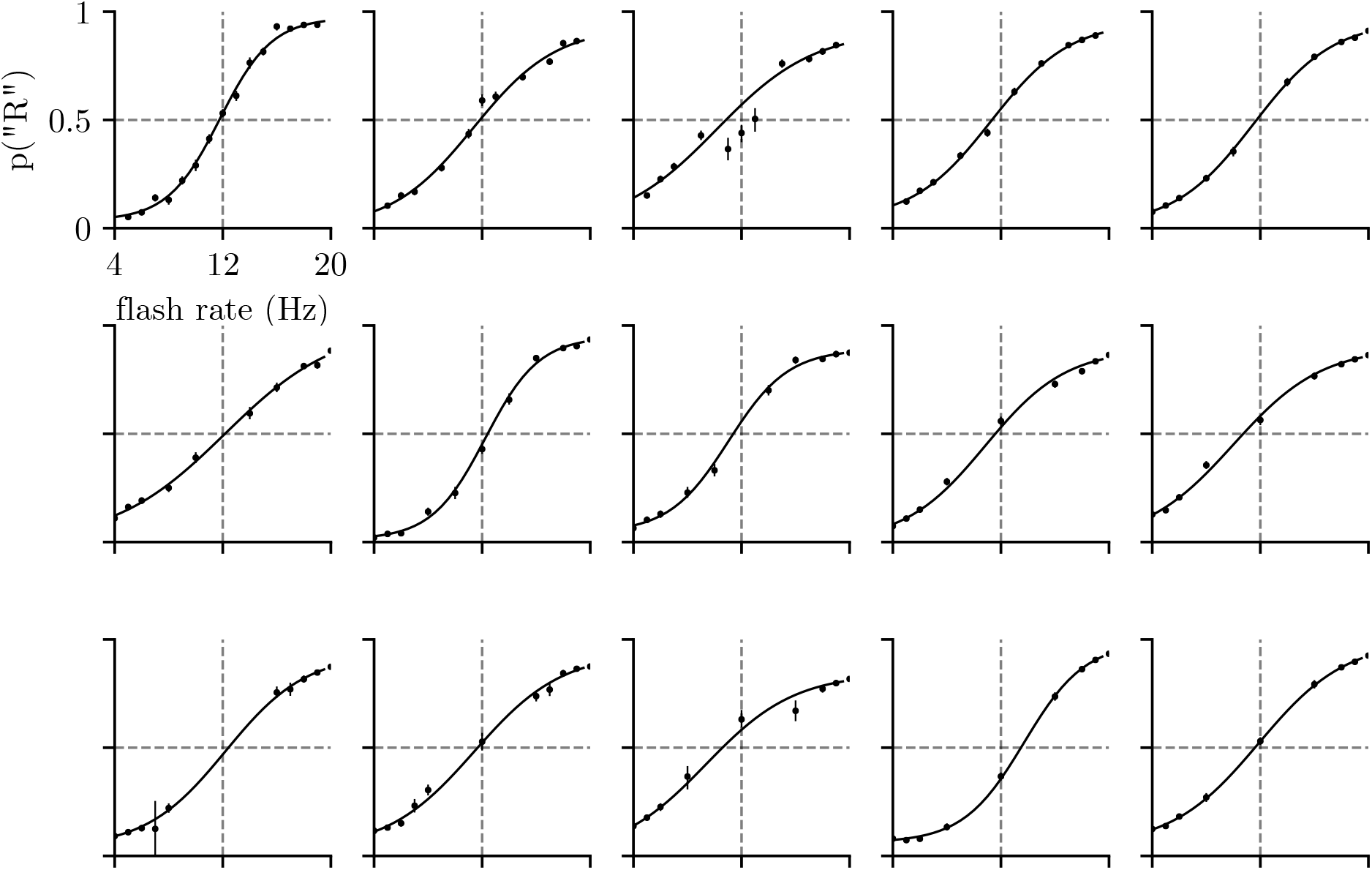
Psychometric curves for Odoemene et al. [20] animals. We plot the psychometric curves for each of the 15 animals whose choice data we study. We also show each animal’s empirical choice probabilities; error bars are 68% confidence intervals (median n across animal-stimulus pairs was n=1070; minimum was n=8; maximum was n=5253). Animals are ordered as in Fig. S13.

